# Avidity-driven polarity establishment via multivalent lipid-GTPase module interactions

**DOI:** 10.1101/303024

**Authors:** Julien Meca, Aurélie Massoni-Laporte, Elodie Sartorel, Denis Martinez, Antoine Loquet, Birgit Habenstein, Derek McCusker

## Abstract

While Rho GTPases are indispensible regulators of cellular polarity, the mechanisms underlying their anisotropic activation at membranes have been elusive. Using the budding yeast Cdc42 GTPase module, which includes a Guanine nucleotide Exchange Factor (GEF) Cdc24 and the scaffold Bem1, we find that avidity generated via multivalent anionic lipid interactions is a critical mechanistic constituent of polarity establishment. We identify Cationic-enriched Lipid Interacting Clusters (CLICs) in Bem1 that drive the interaction of the scaffold-GEF complex with anionic lipids at the cell pole. This interaction increases lipid acyl chain ordering, thus contributing to membrane rigidity and feedback between Cdc42 and the membrane environment. Sequential mutation of the Bem1 CLIC motifs, PX domain and the PH domain of Cdc24 lead to a progressive loss of cellular polarity stemming from defective Cdc42 nanoclustering on the plasma membrane and perturbed signaling. Our work demonstrates the importance of avidity via multivalent anionic lipid interactions in the spatial control of GTPase activation.

## Introduction

Cellular polarity, the anisotropic organization of cellular constituents, is essential for basic cellular functions including migration, division and polarized growth ^1^. In diverse eukaryotes, cellular polarity is regulated by GTPases of the Rho family, including Cdc42 ^2–6^. The temporal and spatial control of Cdc42 activity on the plasma membrane ensures the anisotropic activation of the protein, and thus its function as an essential regulator of cellular polarity.

Cdc42 is prenylated at its C-terminus via the covalent addition of an S-geranylgeranyl group and methylesterification of a cysteine residue ^7^. These modifications, together with juxtaposed polybasic residues, facilitate the high affinity binding of the GTPase to membrane ^8, 9^. While lipid modification ensures the membrane association of the protein, it is not sufficient to account for its anisotropic localization ^10, 11^. Rather, the enrichment of Cdc42 at the cell pole is thought to reflect the local activation of the GTPase by its GEF Cdc24 and stabilization of the GEF-Cdc42 complex involving the scaffold Bem1 ^12, 13^

How then, are Bem1 and Cdc24 recruited to the plasma membrane to locally activate Cdc42? Both are peripheral membrane proteins containing a Pleckstrin-Homology (PH) domain in the case of Cdc24 and a Phox (PX) domain in Bem1. However, the PH domain in Cdc24 displays no detectable phosphoinositide specificity *in vitro*, nor is fusion of the PH domain to GFP sufficient for membrane targeting ^14^ These features reflect the low affinity of the Cdc24 PH domain for phosphoinositides (PI) (K_d_ > 20 μM for PI4,5P2, for example)^14^. The PX domain of Bem1 binds PI3P, PI4P and phosphatidylserine (PS) *in vitro*. While the affinity of the Bem1 PX domain for PI3P is low (K_d_ > 100 μM), its interaction with PS and PI4P was observed to be stronger ^15, 16^ However, neutralization of a critical arginine required for electrostatic interactions with anionic lipids within the Bem1 PX domain did not disrupt Bem1 localization unless additional pathways that guide Cdc42 activation were also blocked ^17^ The mechanism accounting for the site-specific recruitment of these critical Cdc42 regulators to the plasma membrane therefore represents a key unanswered question.

At the plasma membrane of diverse eukaryotes, anionic lipids including PS, contribute to the net negative charge of the membrane and the recruitment of peripheral membrane proteins ^18, 19^ Moreover, the localization of a PS reporter is anisotropic with respect to both the lateral plane of the plasma membrane and the inner leaflet of the membrane ^18^ PS has been shown to play an important role in the anisotropic localization of Cdc42 and Bem1 in *Saccharomyces cerevisiae* ^20, 21^. Work from our lab indicates that PS is required for the spatial organization of Cdc42 in nanoclusters on the plasma membrane, whose size correlates with Cdc42 activity. The addition of PS to wild type cells results in larger Cdc42 nanoclusters at the cell pole in a Bem1-dependent manner ^22^. PS is emerging as a key regulator of nanoclustering in diverse signaling pathways, including Ras signaling ^23–25^ Defining how Cdc42 activators interact with anionic lipids including PS would therefore address one of the most upstream events during polarity establishment and may provide insight into more general features of Ras-family signaling that are conserved among eukaryotes.

Here, using complementary quantitative *in vivo* imaging and reconstitution experiments, we identify a mechanism through which the Cdc42 regulators are recruited to anionic lipids. The interaction of Bem1 with PS and PI4P induces lipid ordering and consequent membrane rigidity. The affinity of Bem1 for anionic lipids is also sufficient to target the associated GEF to this membrane environment, while robust membrane targeting of the GEF-scaffold complex involves multivalent protein-lipid interactions that promote Cdc42 nanoclustering, signaling and cell polarity.

## Results

### PI4P and PS are essential for plasma membrane targeting of Cdc42 regulators *in vivo*

The sole PS synthase *CHO1* is not an essential gene in budding yeast, whereas the anisotropic localization of Cdc42 is essential, suggesting that other lipids may compensate for the lack of PS in *cho1Δ* cells to promote Cdc42 polarization. Recent work uncovered a mechanism of PS transport from the ER to the plasma membrane involving the counter transport of PI4P in the opposite direction ^26, 27^. Thus, it is possible that plasma membrane PI4P levels may be elevated in *cho1Δ* cells, prompting us to examine whether this PI4P pool could account for the residual Cdc42 localization in the *cho1Δ* mutant.

First, we examined the localization of Bem1-GFP in *cho1Δ* cells and confirmed that the percentage of cells displaying polarized Bem1-GFP was reduced compared to a wild type (WT) control population (26% cells with polarized Bem1-GFP in *cho1Δ*, 71% in WT; Fig. 1a, b)^21^. In addition, the fluorescence intensity of Bem1-GFP was also reduced at the pole of those *cho1Δ* cells that displayed polarized Bem1-GFP (Supplementary Fig. 1a). We reasoned that the residual cohort of *cho1Δ* cells might display polarized Bem1-GFP due to elevated PI4P levels, since PI4P would be predicted to accumulate at the plasma membrane in the *cho1Δ* mutant as a consequence of reduced PI4P-PS exchange. Consistent with this reasoning, and with the previously reported elevated global PI levels in *cho1Δ* cells detected by mass spectrometry ^21^, we observed an approximately 2.5-fold increase in the intensity of a PI4P probe by quantitative imaging in *cho1Δ* cells (Fig. 1c, d). Next, the levels of plasma membrane PI4P and PS were ablated by appending an auxin-inducible degron to the PI-4-kinase Stt4 in the *cho1Δ* mutant ^28, 29^. As expected, the double mutant was sensitive to the presence of auxin and choline in the media (Supplementary Fig. 1b, c). Quantitative imaging indicated reduced plasma membrane levels of a PI4P probe upon auxin treatment (Supplementary Fig. 1d, e). In addition, the levels of 9XMyc-AID-stt4 were reduced after auxin treatment, consistent with its degradation (Supplementary Fig. 1f). The percentage of cells displaying polarized Bem1-GFP when PI4P and PS were ablated dropped from 25% to 3%. Upon PS and PI4P attenuation, Bem1-GFP accumulated in puncta in 55% of cells (Fig. 1e and f). Simultaneous imaging of the GEF Cdc24-mCherry and Bem1-GFP indicated that the Bem1-GFP puncta also contained the GEF, reflecting association of the two proteins in a protein complex (Fig. 1g and h). These results underscore the key role of PS and plasma membrane PI4P in the localization of Bem1 and Cdc24 *in vivo*, and the importance of these anionic lipids in the spatial control of Cdc42 activity.

**Fig. 1.**
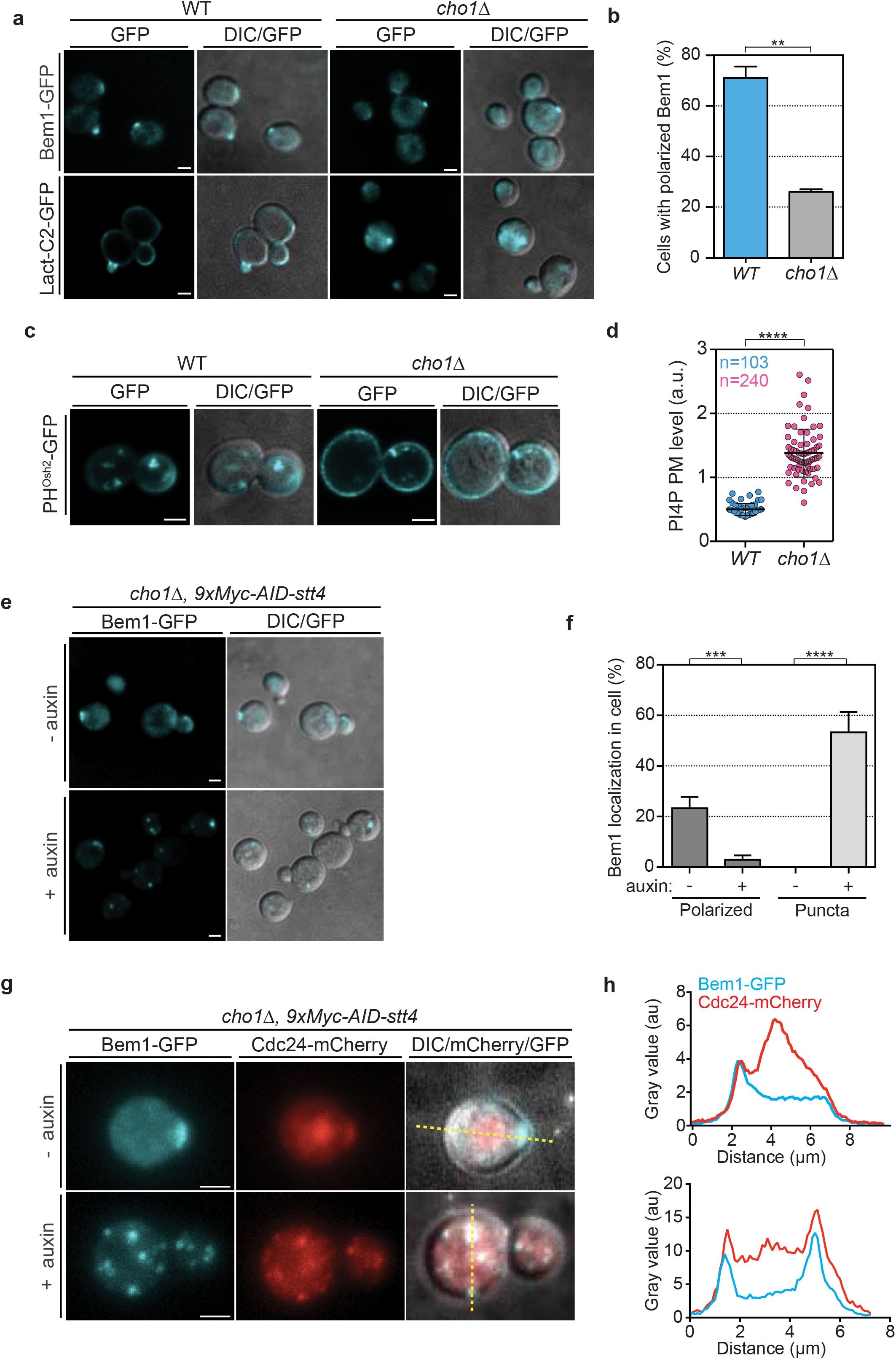
PI4P and PS are essential for the anisotropic plasma membrane targeting of the Cdc42 GEF-scaffold complex *in vivo*. **a**, Representative images of Bem1-GFP and the PS marker LactC2-GFP (cyan) in wild type and *cho1Δ* cells. Merged DIC-fluorescence images are also shown. Images show average intensity projections of deconvolved z-stacks. Scale bars, 2 μm in all images. **b**, Frequency of cells displaying polarized Bem1-GFP signal in wild type and *cho1Δ* cells (n>100 cells counted in each of 3 independent experiments). Bars represent mean and SD. Student t-tests were performed where confidence is **p<0.01). **b**, Imaging of the PI4P probe GFP-2xPH^Osh2^ in wild type and *cho1Δ* cells. Images show average intensity projections of deconvolved z-stacks. **d**, Scatter dot plot showing PI4P levels at the plasma membrane (see experimental procedure for details of the quantification) in wild type and *cho1Δ* cells (n= around 100 cells observed over 3 experiments). Bars represent mean and SD. Mann-Whitney tests were performed where confidence is ****p<0.0001). **e**, Images of Bem1-GFP (cyan) in *cho1Δ 9xMyc-AID-stt4* cells after 30 min treatment with or without 0.5 mM auxin. Merged DIC-fluorescence images are also shown. Images are average intensity projections of deconvolved z-stacks. **f**, Frequency of cells with polarized Bem1-GFP signal or Bem1-GFP in puncta in *cho1Δ 9xMyc-AID-stt4* cells treated with or without auxin as shown in E (n >100 cells counted in each of 6 independent experiments). Bars represent mean and SD. Student t-tests were performed where confidence is ***p<0.001 and ****p<0.0001). **g**, Images of Bem1-GFP (cyan) and Cdc24-mCherry (red) signals in *cho1Δ 9xMyc-AID-stt4* cells with or without auxin as shown in E. Merged DIC-fluorescence images are also shown. Images show maximum intensity projections of z-stacks. **h**, Graphs showing the line-scans (yellow dashed line in G) of Bem1-GFP and Cdc24-mCherry signals. The top and bottom graphs correspond to the top and bottom images in G, respectively. The line-scan reveals the colocalization of Bem1 and Cdc24 at the pole in a non-treated cell and in the puncta in cells treated with auxin.

### Identification of a robust anionic lipid targeting sequence in Bem1

Previously, the PX domain of Bem1 was shown to interact directly with anionic lipids; however, it is unknown whether the interaction is sufficiently robust to target the full-length protein to these lipids ^15, 16^. We established liposome floatation assays to address this question. In the assay, liposomes were mixed with Bem1 purified from bacteria and floated through dense sucrose. In the event of a sufficiently strong interaction, the protein is found in the supernatant, associated with the liposomes (Fig. 2a). A panel of neutral to anionic lipids were tested. While BSA did not interact appreciably with any of the lipids (Supplementary Fig. 2a), Bem1 displayed a robust interaction with anionic liposomes. Strikingly, liposomes containing 20% PS, 5% PI4P and 75 *%* PC, mimicking the composition of the plasma membrane (PM lipids), recruited around 93% of the Bem1 in the assay, indicating that the full-length protein binds strongly to these lipids in the absence of additional proteins (Fig. 2b).

**Fig. 2.**
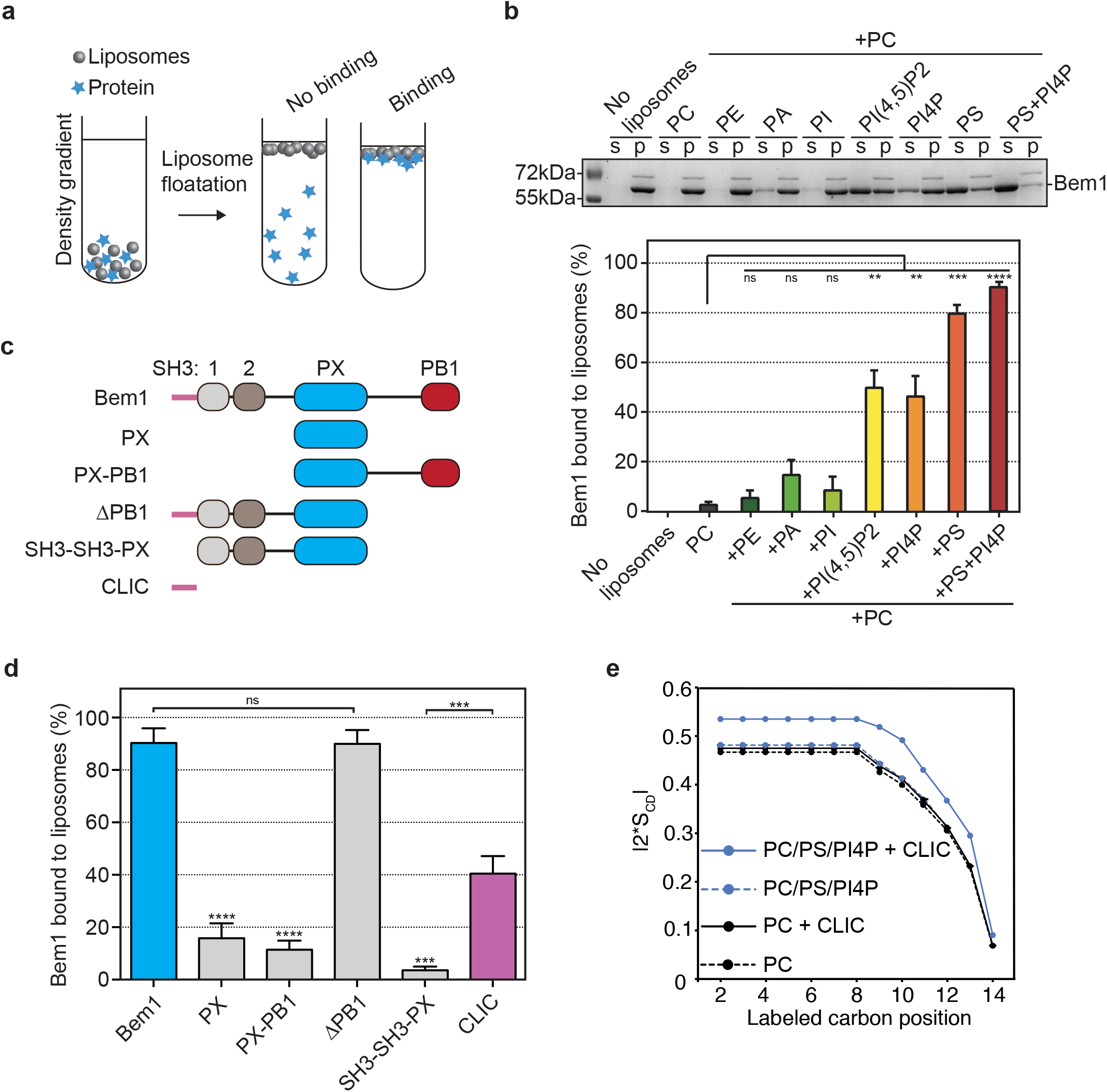
Identification of an anionic lipid targeting sequence in Bem1 and its effect on the ordering of lipid acyl chains. **a**, Schematic of the liposome floatation assay. In the assay, liposomes of defined lipid composition were floated through a dense sucrose gradient by ultracentrifugation. Protein that associates with the liposomes become enriched in the supernatant. **b**, Upper panel. SDS-PAGE stained with Coomassie blue in which Bem1 is indicated. Lower panel. The liposomes were composed of 100% phosphatidylcholine (PC), 80% PC and 20% phosphatidylethanolamine (PE), 95% PC and 5% phosphatidic acid (PA), 95% PC and 5% phosphoatidylinositol (PI), 95% PC and 5% PI(4,5)P2 (PI(4,5)P2), 95% PC and 5% PI4P (PI4P), 80% PC and 20% phosphatidylserine (PS) or 75% PC 20% PS 5% PI4P (PS+PI4P). (S) supernatant, (P) pellet, Lower panel displays the percentage of Bem1 associated with liposomes containing the indicated lipids from 3 independent experiments. Error bars display SD. Student t-tests were performed where confidence is **p<0.01, ***p<0.001 and ****p<0.0001. **c**, Scheme of full-length Bem1 with its domains and the Bem1 deletion constructs used to identify the anionic lipid interacting sequences. **d**, Percentage of the indicated bem1 constructs associated with liposomes containing 75% PC, 20% PS and 5% PI4P. Error bars show SD (n=3 experimental replicates). Student t-tests were performed where confidence is ***p<0.001. **e**, Lipid ordering determined by ^2^H solid-state NMR analysis of liposomes containing PC-d54/PS/PI4P (15:4:1 molar ratio) in the presence or the absence of the Bem1 CLIC motifs. Calculation of oriented-like spectra from Pake patterns (de-Pake-ing) and simulation of ^2^H solid-state NMR spectra were applied to measure individual quadrupolar splittings for PC-d54 and determine order parameter accurately.

The region of Bem1 responsible for the PS-PI4P lipid interaction was next mapped (Fig. 2c). While the PX domain interacted weakly with PM lipids in the assay (16% floatation), an N-terminal 72 amino acid sequence interacted more strongly with this lipid mix (41% floatation)(Fig. 2d, mauve bar). The sequence was enriched in clusters of basic residues that we refer to as a Cationic-enriched Lipid Interacting Clusters (CLICs). We next addressed the mechanism through which the CLICs interact with PM lipids by mixing liposomes with the CLICs and analyzing the interaction using ^2^H solid-state NMR spectroscopy. Strikingly, the CLICs had a structural impact on the entire length of the lipid acyl chain, increasing the degree of order in the 14 carbon atoms of the lipid PC acyl chain in the presence of PS and PI4P, while the acyl chains in liposomes containing only PC remained unaffected by the CLICs (Fig. 2e). These results indicate that Bem1 interacts with anionic lipids via anionic lipid interacting motifs, and in doing so, Bem1 in turn increases ordering along the length of the acyl chain backbone in a reciprocal fashion.

### The Bem1 CLIC motifs can act as a heterologous plasma membrane targeting signal *in vivo*

The N-terminus of Bem1 contains 3 clusters of basic residues, or CLIC motifs, totaling 14 lysine and arginine residues. We mutated each cluster individually or all 14 residues simultaneously (Fig. 3a). We first performed liposome floatation assays to identify the CLIC motif that contributed most to Bem1 anionic lipid targeting. Of the 3 clusters of basic residues, the most N-terminal CLIC-1 cluster appeared to be the most important (Fig. 3b). In addition, mutation of all basic residues to alanine (*clic*-14A) or a charge swap to glutamate (*clic*-14E) strongly attenuated the interaction of these full-length Bem1 constructs with PM lipids *in vitro*. We next verified that the resulting clic-14E mutant protein retained the ability to boost Cdc24 GEF activity in a GEF assay, and had thus not been non-specifically damaged by the charge-swap mutations. This FRET-based mant-GTP loading assay serves as a sensitive readout for Bem1 function, since Bem1 boosts GEF activity via interactions with Cdc24 and Cdc42 that are distinct from the N-terminal CLIC motifs ^30–32^ The clic-14E mutant was chosen for these experiments because it was expressed at similar levels to wild type Bem1 *in vivo*, as demonstrated below. In these assays, the bem1 clic-14E mutant and wild type Bem1 boosted Cdc24 GEF activity indistinguishably, consistent with the idea that the clic mutations had not resulted in non-specific Bem1 mis-folding, (Fig. 3c and Supplementary Fig. 2b).

**Fig. 3.**
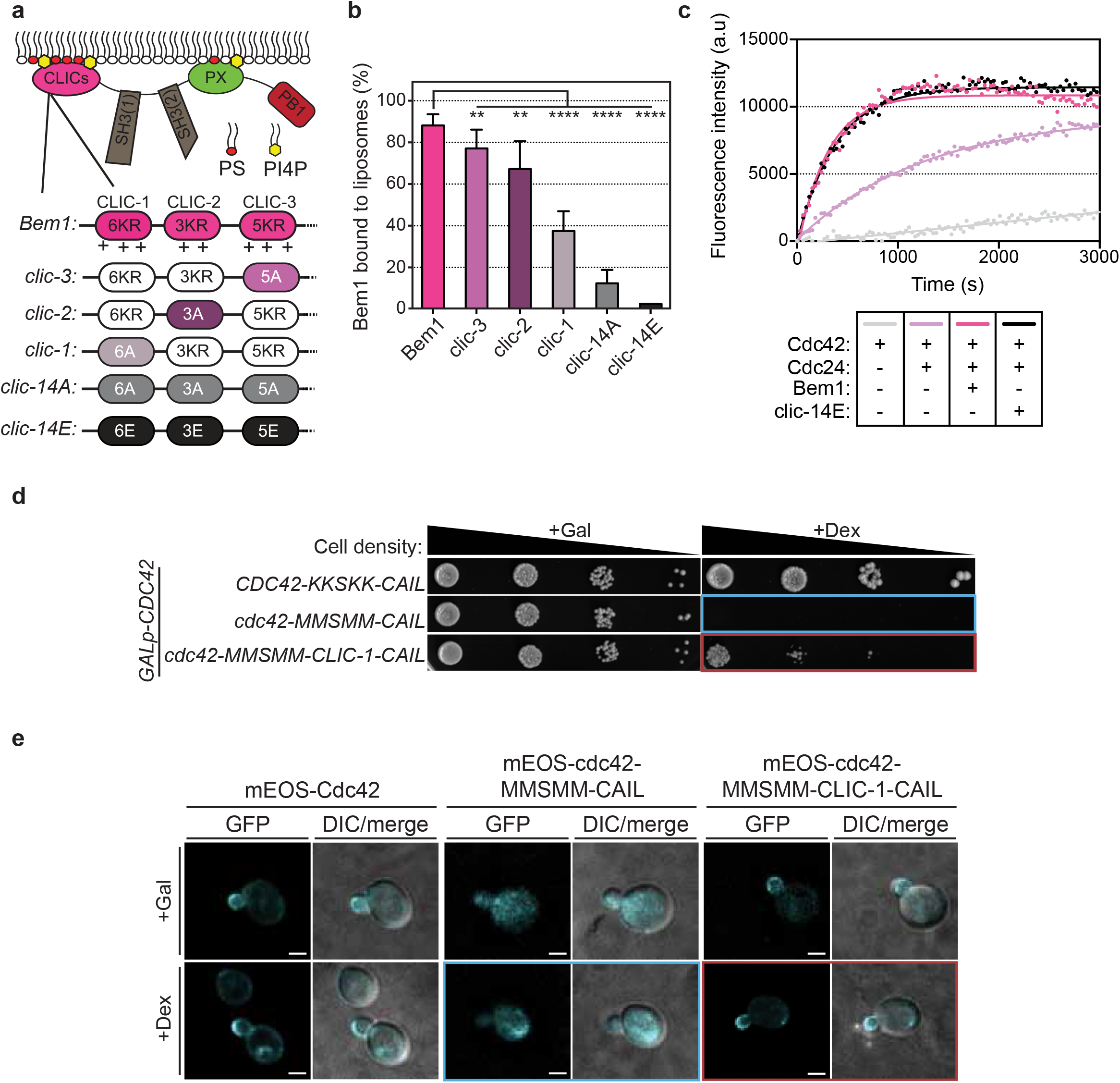
The Bem1 CLIC sequence can act as a heterologous plasma membrane targeting signal *in vivo*. **a**, Schematic showing Bem1 domains with the lipid interacting motifs including the cationic lipid interacting clusters (CLICs) and the PX domain. The Bem1 N-terminus contains 3 CLICs. The first cluster (CLIC-1) is composed of 6 K/R residues, the second cluster (CLIC-2) of 3 K/R residues and the third cluster (CLIC-3) of 5 K/R residues. The scheme shows the full-length bem1 constructs where none, one or all CLICs were mutated. **b**, Percentage of the different full-length bem1 clic mutants associated with liposomes containing 75% PC 20% PS and 5% PI4P. Error bars display SD (n=3 experimental replicates). Student t-tests were performed where confidence is **p<0.01 and ****p<0.0001. **c**, Fluorescence intensity change associated with the nucleotide exchange of GDP-Cdc42 for mant-GTP Cdc42. Fluorescence was measured after the addition of GDP-Cdc42 to reactions containing Mant-GTP (100 nM), GMP-PNP (100 μM) and the proteins indicated. **d**, Ten-fold serial dilutions of cells and subsequent colony formation on the indicated plates, where expression of wild type *GALp-CDC42* is either induced in the presence of Gal or repressed in Dex. Note how mutation of the wild type Cdc42 (KKSKK) to MMSMM is lethal (see blue box), whereas appending the Bem1 CLIC-1 motif to this *cdc42* mutant restores viability (red box). **e**, Representative images of the mEOS-cdc42 mutants signal (cyan) after inducing the expression of *GAL1p-CDC42* in the presence of Gal or repressing it in the presence of Dex. The cells in the blue and red boxes correspond to the cells in the blue and red box in panel **d**. Images are average fluorescence intensity projections. Merged DIC-fluorescence images are also shown.

The CLIC motifs in Bem1 are necessary for the interaction with anionic lipids, but are they sufficient? We tested whether a single CLIC motif in Bem1 was sufficient for heterologous targeting of proteins to the plasma membrane *in vivo*. The viability of budding yeast requires a Cdc42 C-terminal polybasic sequence containing 4 lysines that interact with anionic lipids, providing an *in vivo* system in which to test the functional importance of a single CLIC motif ^9^ We generated budding yeast in which the wild type copy of *CDC42* was expressed from the conditional *GAL1* promoter. Expression of wild type *CDC42* was repressed by plating cells on dextrose and then we tested whether the most N-terminal Bem1 CLIC-1 motif could support the viability of the *cdc42* polybasic mutant when engineered onto the C-terminus of cdc42 immediately preceding the geranylgeranylation site. While the *cdc42* polybasic mutant was unable to support cellular viability when grown on dextrose, as indicated by the lack of colony growth on dextrose plates (Fig. 3d, blue box), appending the Bem1 CLIC-1 motif to this mutant restored viability, albeit with a reduced rate of colony formation compared to wild type control cells (Fig. 3d, red box). Control experiments indicated that the loss of *CDC42* function required mutation of all 4 C-terminal lysines (Supplementary Fig. 2c). The CLIC-1 motif was also sufficient to target the polybasic *cdc42* mutant to the plasma membrane in an anisotropic manner (Fig. 3e, red box). These results indicate that the CLIC motifs are necessary for the strong interaction between Bem1 and anionic lipids in the reconstituted assay, and that the affinity of the single CLIC-1 motif for these lipids is sufficient to support viability when heterologously appended to a *cdc42* lipid-binding mutant *in vivo*.

### The Bem1 CLIC sequence is required for Bem1 targeting to the cell pole *in vivo*

We next addressed the importance of the anionic lipid targeting CLIC motifs within Bem1 *in* vivo. Since previous work demonstrated that the PX domain in Bem1 was only important for the localization of the protein when additional pathways that guide polarity were also inactivated ^17^, we reasoned that the PX domain may function with the CLIC motifs in a multivalent fashion to confer robust membrane targeting. The importance of the CLIC and PX membrane targeting sequences were tested *in vivo* by replacing the wild type copy of *BEM1* with the *bem1 clic-14E* mutant, a *bem1 px* domain mutant (K338M, K348A, R349A, R369A)^15^, or a mutant in which both sequences were mutated. As a measure of Bem1 function, the rate of colony formation at 37ºC was observed, since *bem1Δ* cells display a growth defect at this temperature that has been attributed to defective organization of the actin cytoskeleton ^33, 34^ While the *bem1 px* mutant displayed a rate of colony formation indistinguishable from wild type cells, the *bem1 clic-14E* mutant displayed a reduced rate, which was exacerbated in the double *bem1-14E px* mutant (Fig. 4a). The mutant proteins were not non-specifically destabilized by the mutations, since 3xHA-tagged mutants were expressed comparably to wild type Bem1-3xHA (Fig. 4b).

**Fig. 4.**
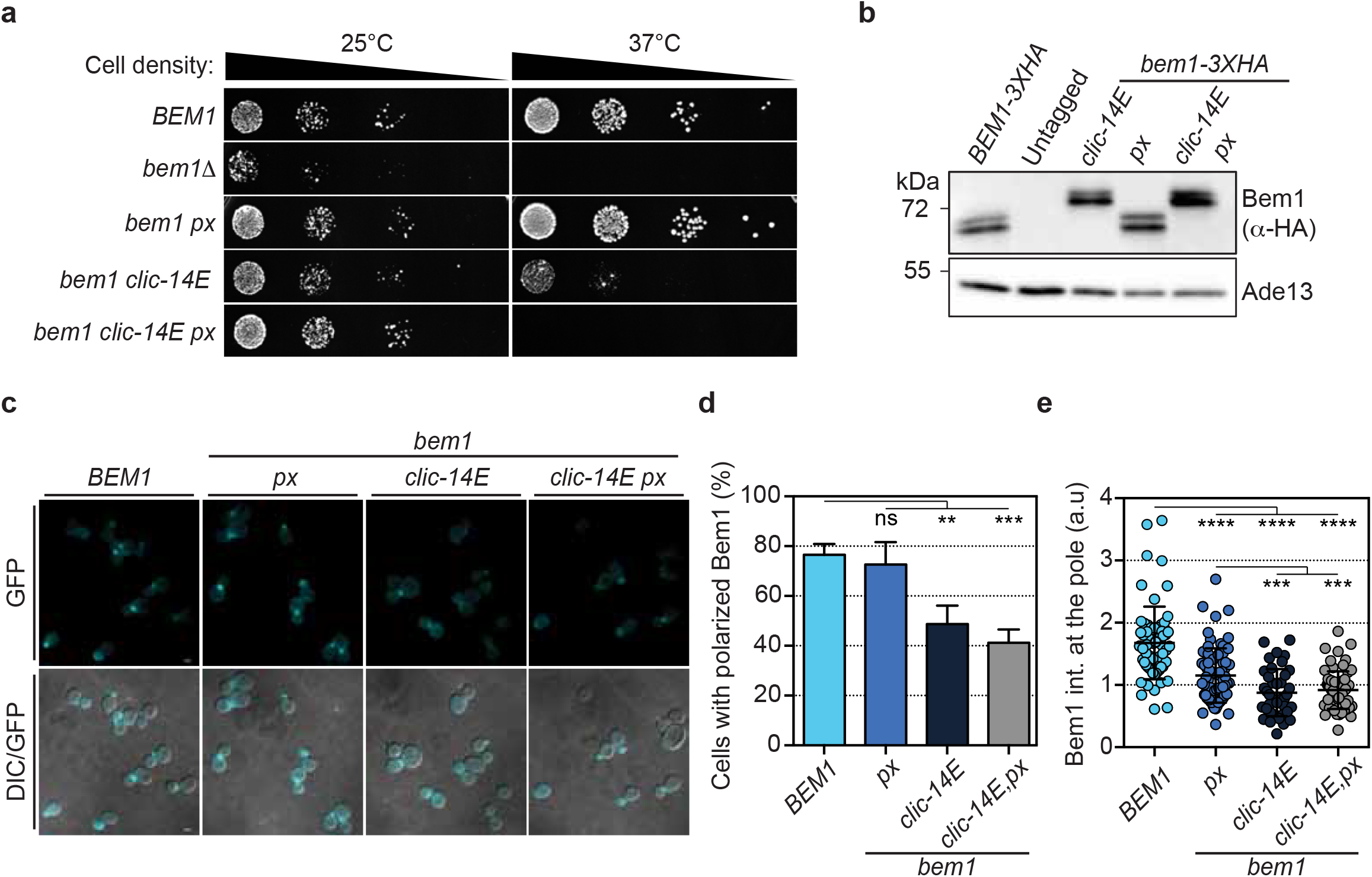
The Bem1 CLIC sequence is required for Bem1 targeting to the cell pole *in vivo*. **a**, Ten-fold serial dilutions of the indicated mutant cells and subsequent colony formation at the temperatures indicated. Note how mutations on both the CLICs and PX in bem1 compromise growth at the restrictive temperature. **b**, Western blots probed with anti-HA antibody to detect the indicated bem1 constructs tagged with 3XHA (top panel). Detection of Ade13 was used as loading control (bottom panel). **c**, Representative images of the indicated bem1-GFP construct signals (cyan, top panels). The GFP images are average intensity projections. GFP images were also merged with DIC (bottom panels). **d**, Frequency of cells with polarized Bem1-GFP fluorescence (n>100 cells in each of at least 3 independent experiments). Bars display the mean and SD. Student t-tests were performed where confidence is **p<0.01 and ***p<0.001). **e**, Scatter dot plot showing the level of the indicated bem1-GFP construct at the cell pole (n>100 cells observed over 3 experiments). Bars indicate mean and SD. Mann-Whitney tests were performed where confidence is ***p<0.001 and ****p<0.0001).

The percentage of cells displaying polarized GFP fluorescence was reduced in the *bem1 clic-14E-GFP* mutant compared to wild type, and even more diminished in the *bem1 *clic*-14E px-GFP* double mutant (77% wild type, 49% **clic*-14E* and 41% **clic*-14E px* double mutant, Fig. 4c, d). Moreover, the intensity of GFP fluorescence at the pole in single cells indicated that all three mutants displayed significantly reduced levels of GFP signal compared to wild type cells (Fig. 4e). These results, which are consistent with our reconstitution experiments, identify the importance of the CLIC motifs in targeting Bem1 to the cell pole *in vivo*.

### Multivalent protein-lipid interactions drive avid targeting of the GTPase module to anionic lipids *in vitro* and *in vivo*

Previous work reported that the GEF Cdc24 is targeted to, but not maintained at the pole in *bem1Δ* mutants ^12^ Conversely, Bem1 polarization is not maintained in *cdc24* mutants ^12^, suggesting that Cdc24 may display some affinity for lipids *in vivo*, likely via its PH domain ^14^ This led us to test whether multivalent protein-lipid interactions in Bem1 and Cdc24 may drive recruitment of the GEF-scaffold complex to the pole (Fig. 5a). First, we tested whether Cdc24 has appreciable affinity for anionic lipids in the liposome floatation assay. Approximately 33% of Cdc24 interacted with PM lipids in the assay, and the association was reduced to around 15% when a mutation was introduced into a conserved cationic residue in the beta-2 sheet of the PH domain (cdc24 K513A, which we refer to as cdc24 ph)(Fig. 5b and Supplementary Fig. 2e)^35^. The addition of Bem1 dramatically increased the amount of Cdc24 associated with PM lipids from 33% to more than 88% (Fig. 5c). While mutation of the CLIC motifs in Bem1 had the strongest impact on Cdc24 association with PM lipids, successive neutralization of the CLIC and PX motifs in Bem1, combined with mutation of the PH domain in Cdc24 resulted in a progressive reduction in the interaction of Cdc24 with PM lipids (Fig. 5c). We directly tested whether avidity is generated via multivalent interactions by varying the concentration of Bem1 and plotting the amount of Cdc24 associated with PM lipids (Fig. 5d). Fitting the curves with an equilibrium-binding model revealed that the K_d_ of Cdc24 for anionic lipids was acutely sensitive to Bem1 lipid binding. The apparent K_d_ of Cdc24 for PM lipids in the presence of wild type Bem1 was 6 nM, 117 nM in the bem1 clic-14A mutant and 180 nM in the bem1 clic-14A px mutant. These results indicate that multivalent lipid binding motifs in Bem1, conferred by the CLIC motifs and PX domain, contribute to the avid targeting of Cdc24 GEF activity to anionic lipids in the reconstituted system. Of these interactions, the CLIC motifs that we identify in Bem1 provide the strongest anionic lipid targeting to the GEF-scaffold complex. These results are consistent with a model of GEF-scaffold targeting via multivalent anionic lipid avidity.

**Fig. 5.**
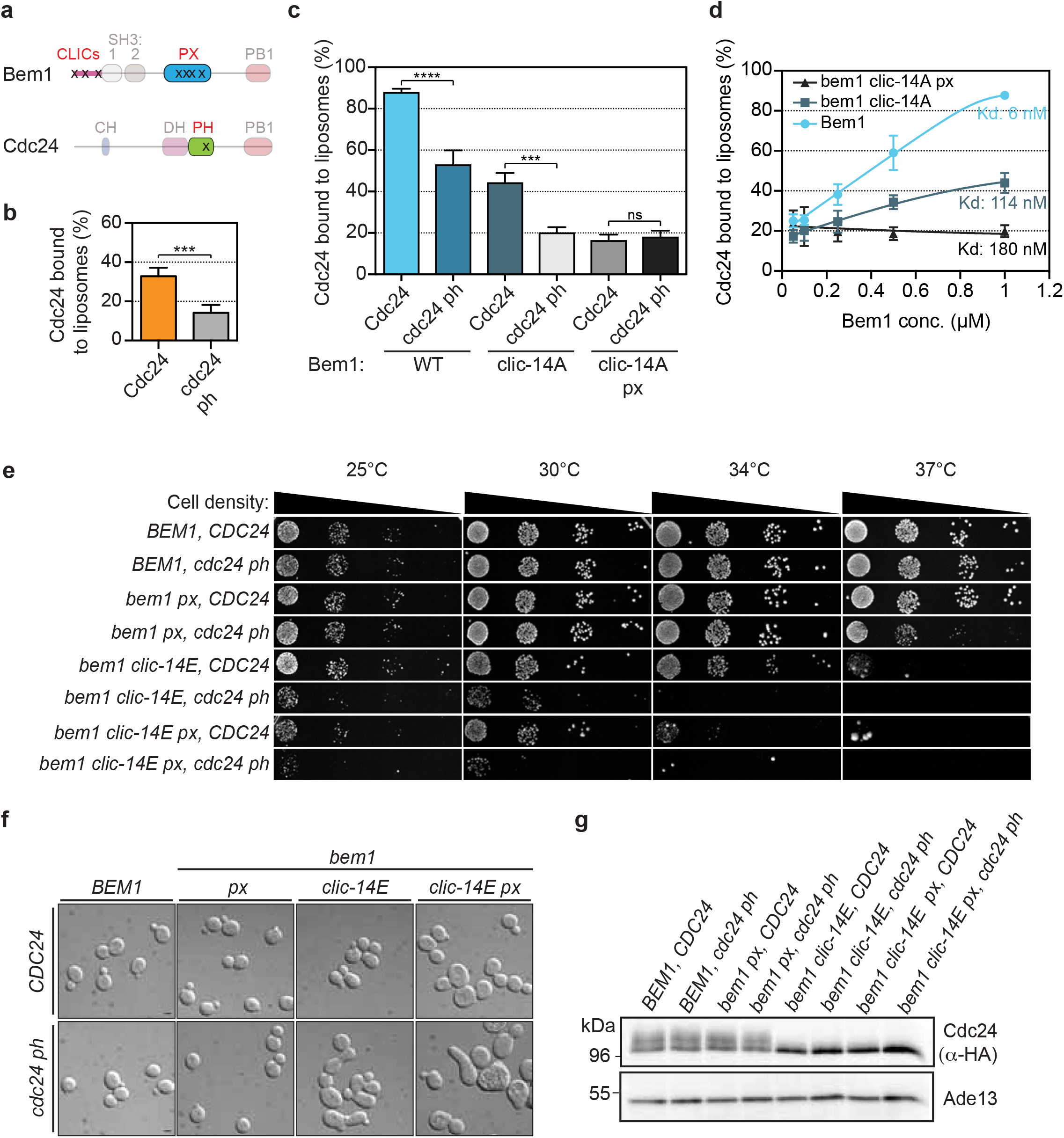
Multivalent protein-lipid interactions drive avid targeting of the Bem1-Cdc24 complex to anionic lipids. **a**, Scheme of Bem1 and Cdc24 proteins indicating the relative position of the mutations in the lipid tethering motifs (black x). **b**, Percentage of full-length Cdc24 and cdc24 ph domain mutant (cdc24 ph) associated with liposomes containing 75% PC 20% PS and 5% PI4P. Error bars display SD (n=3 experimental replicates). Student t-tests were performed where confidence is ***p<0.001. **c**, Percentage of Cdc24 and cdc24 ph mutant associated with liposomes of the composition shown in B in the presence of the indicated bem1 protein. Note how additive mutations in the Bem1 lipid binding sequences reduce the percentage of Cdc24 associated with the liposomes. **d**, Percentage of Cdc24 and cdc24 ph associated with liposomes of the composition shown in B as a function of the indicated bem1 protein concentration. The curves denote the regression fit of the data to equation 2 in the materials and methods. Error bars correspond to SD (n=3 experimental replicates). Student t-tests were performed where confidence is ***p<0.001 and ****p<0.0001. **e**, Ten-fold serial dilutions and subsequent colony formation of the indicated mutant cells at the indicated temperatures. **f**, DIC images of the indicated bem1 and cdc24 mutants showing the increased morphological defects ensuing from loss of lipid tethering in the *bem1* and *cdc24* mutants. **g**, Western blots probed with anti-HA antibody to detect Cdc24 or cdc24 ph tagged with 3XHA in the indicated mutant strains (top panel). Detection of Ade13 was used as a loading control (bottom panel).

The multivalent avidity model was next tested *in vivo*. Mutation of the Bem1 CLICs, PX domain and the Cdc24 PH domain resulted in a progressively more pronounced temperature sensitive phenotype (Fig. 5e). Importantly, by appending a geranylgeranylation sequence to these mutants, it was possible to restore growth at 37ºC, indicating that temperature sensitivity was a result of reduced membrane targeting and not non-specific protein mis-folding due to mutation (Supplementary Fig. 2d). Morphological defects consistent with a loss of cell polarity in the mutants also became more severe as additional mutations in the identified lipid tethering motifs were combined, even at 25ºC (Fig. 5f). We addressed whether the loss of cellular polarity observed *in vivo* was associated with specific Cdc42 signaling pathway defects. Cdc24 multi-site phosphorylation by the p21 Activated Kinase (PAK) Cla4 occurs optimally in the presence of Bem1 when Cdc24 is localized on the plasma membrane ^12, 31, 36, 37^. We therefore predicted that Cdc24 might display aberrant phosphorylation in the GEF-scaffold lipid-binding mutants. Consistently, Cdc24 phosphorylation was observed to be dramatically reduced in the *bem1 clic-14E* mutant, and all combinations thereof, as indicated by the increased hypophosphorylated form of Cdc24 in electrophoretic mobility shifts during SDS-PAGE (Fig. 5g). These results indicate that the CLIC motifs identified in Bem1 and the multivalent anionic lipid interactions displayed by the GEF-scaffold complex are required for the spatial control of Cdc42 activation, signaling via PAK, and the ensuing control of cellular polarity.

### Scaffold tethering to anionic lipids affects Cdc42 dynamics and activation *in vivo*

The loss of cell polarity ensuing from reduced Bem1 tethering to the plasma membrane suggested that Cdc42 dynamics would be altered in the *bem1 clic-14E* mutant. At the cell pole in wild type cells, Cdc42 displays reduced diffusion compared with elsewhere on the plasma membrane, reflecting activation of Cdc42 at the cell pole ^22, 38^. Previous work from our lab demonstrated that Bem1, which boosts Cdc42 activation ^31^, and PS, which recruits Cdc42 activators, contribute to the reduced diffusion and nanoclustering of Cdc42 at the pole ^22^ We therefore reasoned that the reduced rate of Cdc42 diffusion and its nanoclustering at the pole may be linked to the lipid rigidification exerted by the Bem1 CLIC motifs. To test this hypothesis, we monitored mEOS-Cdc42 dynamics in live wild type, *bem1 clic-14E* and *bem1 clci-14E px* mutant cells by single particle tracking Photoactivation Localization Microscopy (sptPALM)(Fig 6a). From high-frequency sptPALM acquisitions (50 Hz), trajectories were obtained from mEOS-Cdc42 at the pole and non-pole regions of cells. The mean square displacement of the protein, which is a measure of Cdc42 mobility, was extracted from the assembled single molecule tracks. In wild type cells, mEOS-Cdc42 displayed more confinement at the pole than at the non-pole, as expected. However, reduced mobility of mEOS-Cdc42 was not observed at the pole of *bem1 clic-14E* and *bem1 clic-14E px* mutants (Fig. 6b). This was also borne out quantitatively by calculating the diffusion coefficient, D, from the slope of the MSD curves (Fig. 6c). We next fixed cells and looked at the organization of mEOS-Cdc42 by PALM. Whereas mEOS-Cdc42 nanoclusters were larger at the pole of wild type cells, we observed no difference in the size of mEOS-Cdc42 nanoclusters between the pole and non-pole of *bem1 clic-14E px* mutant cells (Fig. 6d, e). These results indicate that the interaction between the CLIC motifs in Bem1 and anionic lipids, which rigidifies the lipid acyl chain, is required for the reduced diffusion of Cdc42 and its organization in large nanoclusters at the pole. In order to directly link the alteration in Cdc42 nanoclustering with Bem1 lipid tethering and Cdc42 activation, we monitored Cdc42-GTP levels by quantitative imaging using a gic2(i_-208_)-yeGFP probe, which contains a CRIB motif that interacts with the active GTPase. The levels of the probe were reduced in the *bem1 clic-14E* and *clic-14E px* mutant compared to a wild type control (Fig. 6f). Collectively, these results indicate that Bem1 lipid tethering via the CLIC motifs is required for three key properties of Cdc42 at the cell pole: its reduced diffusion, its organization in large nanoclusters and its optimal activation.

**Fig. 6.**
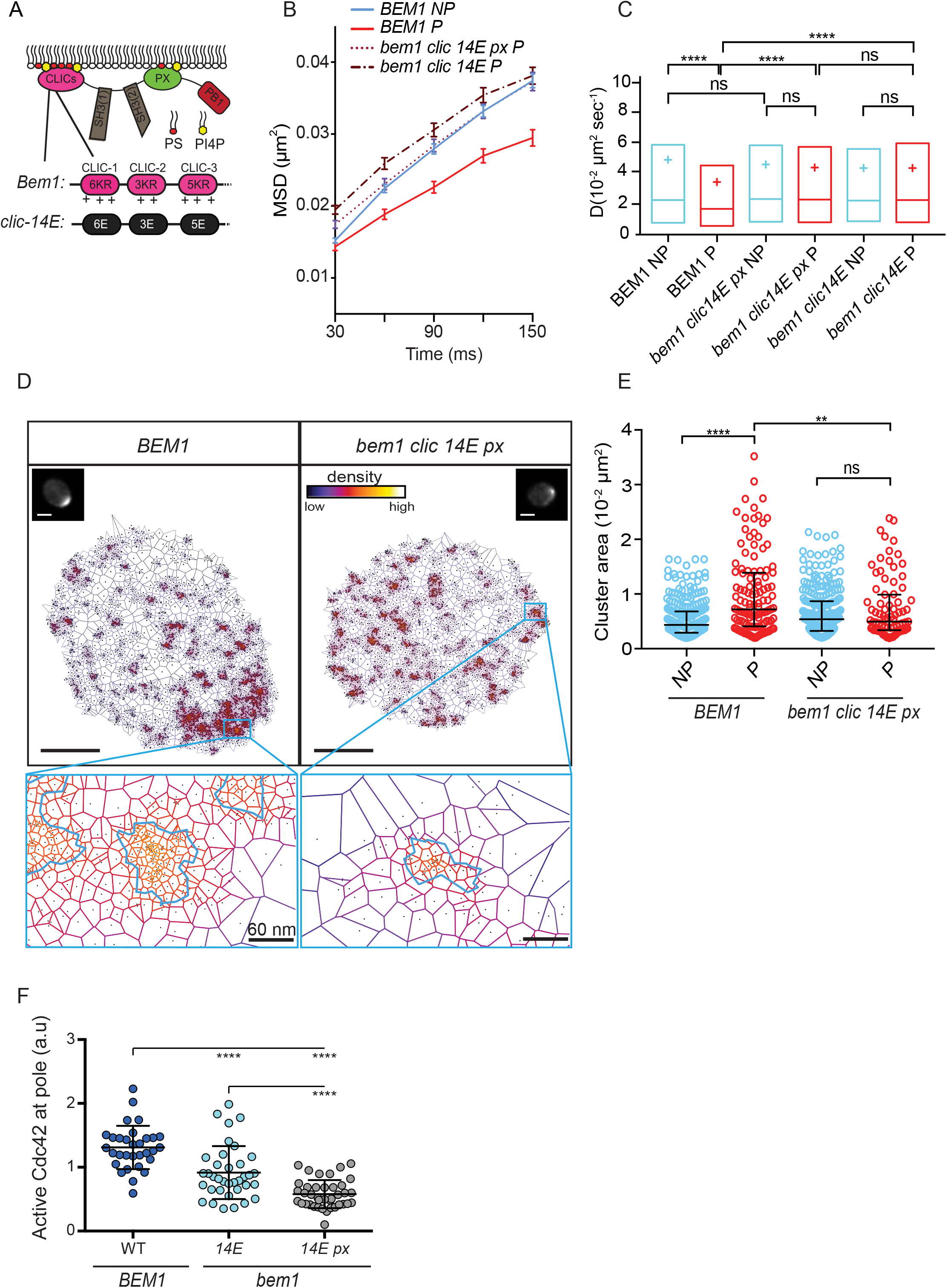
The Bem1 CLIC motifs are required for reduced Cdc42 diffusion, large nanoclusters and optimal Cdc42 activation at the cell pole. **a**, Scheme of the *bem1* mutants used for imaging. **b**, Global average MSD curves of mEOS-Cdc42 in the strains indicated at the Pole (P) and Non-Pole (NP) of cells. Trajectories longer than 6 frames were analyzed. Number of trajectories analyzed: *BEM1* (N=11 cells: NP: 1854 tracks; P: 706); *bem1 clic-14E px* (N=10 cells: NP: 1793 tracks; P: 908); *bem1 clic-14E* (N=13 cells: NP: 1714 tracks; P: 975). **c**, D coefficients of mEOS-Cdc42 in the strains indicated (in box-plots displaying the median (line), the 25-75 percentiles (box) and the mean (cross)), which were compared using a non-parametric, two-tailed Mann-Whitney rank sum test. The resulting P-values are indicated as follows: ns, P > 0.05; *P < 0.05; **P<0.01,***P<0.001,****P<0.0001. **d**, SR-Tesseler images of mEOS-Cdc42 nanocluster organization in *BEM1* (5312 localizations shown in image) and *bem1 clic-14E px* cells (3739 localizations shown). Insets show mEOS-Cdc42 after 491 nm widelfield laser excitation to identify the cell pole. Scale bar: 2 μm. A zoom of the pole region shows the organization of the detected nanoclusters, circled in light blue, in the strains indicated. Scale bar in the zoom: 60 nm. **e**, Distribution of nanocluster area at the pole (P) and non-pole (NP) regions of *BEM1* (diameter NP: 59 nm ± 1 nm (s.e.m); P: 74 nm ± 3,2 nm) (N=16 cells. NP: 570 clusters; P: 162 clusters) and *bem1 clic14E px* cells (diameter NP: 59 nm ± 1,5 nm (s.e.m); P: 57 nm ± 2,4 nm) (N=15 cells. NP: 421 clusters; P: 141 clusters). Data are presented as scatter dot-plots displaying the median as a line and the 25-75 percentiles. Data were compared using non-parametric, two-tailed Mann-Whitney rank sum test. **f**, Active Cdc42-GTP levels were quantified in the strains indicated using a gic2(i_-208_)-yeGFP probe. Bars indicate mean and SD (n<30 cells observed over 2 experiments). Mann-Whitney tests were performed where confidence is ****p<0.0001.

## Discussion

The mechanisms underlying the targeting of the Cdc42 regulators Bem1 and Cdc24 to the plasma membrane represent a longstanding enigma, despite the budding yeast polarity system being one of the most intensively studied among eukaryotes. Previous studies in diverse experimental models have highlighted a crucial role for positive feedback in amplifying the levels of active polarity factors at the cell pole during polarity axis establishment ^39–43^ In budding yeast, Bem1 is proposed to play a role in this feedback ^44–46^; however, the mechanisms that localize Bem1 to the plasma membrane to trigger the positive feedback have been enigmatic. Combining the rapid ablation of plasma membrane lipids *in vivo* with a sensitive reconstituted system and solid-state NMR spectroscopy, we identify a mechanism underlying the spatial tethering of Bem1 and the GEF Cdc24 to anionic lipids enriched at the cell pole. The interaction between Bem1 and anionic lipids is reciprocal in that Bem1 induces the ordering of lipid acyl chains, rigidifying the local membrane environment (Fig. 7). In a wider context, our work also identifies the critical role of multivalent protein-lipid interactions in the control of cellular polarity.

**Fig. 7.**
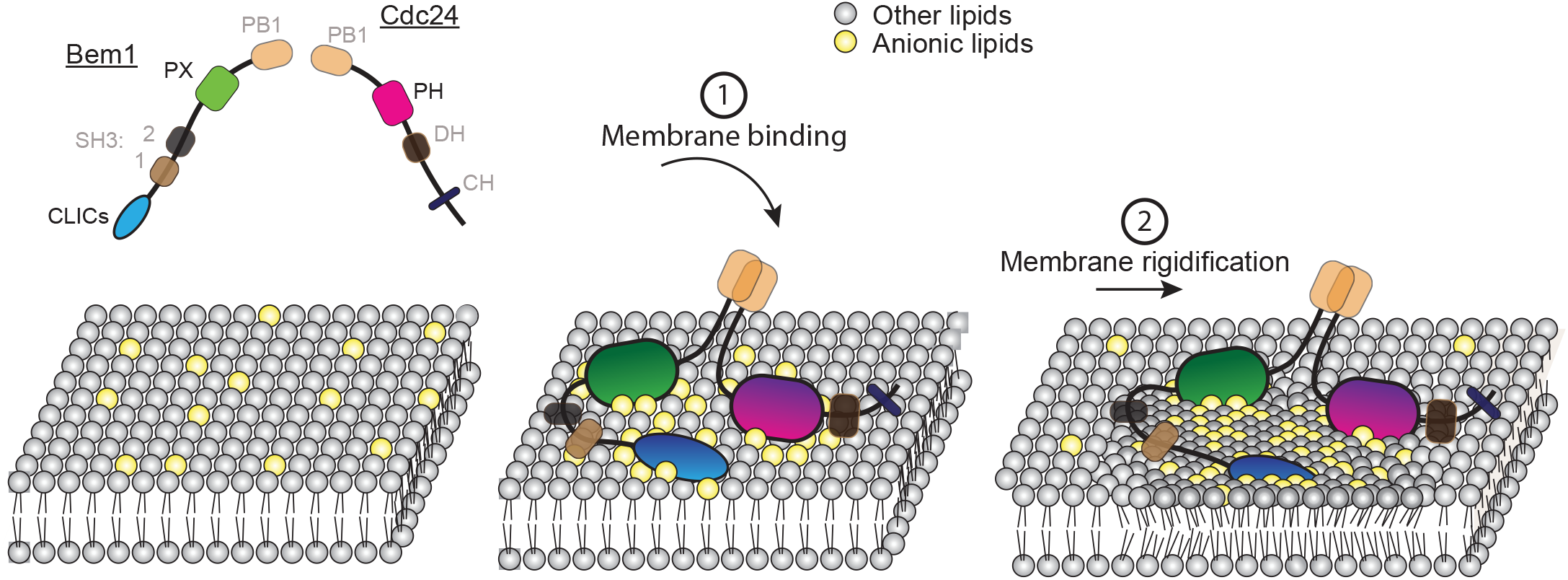
Schematic illustrating the reciprocal relationship between Cdc42 regulators and the membrane environment. 1) The Bem1-Cdc24 complex is recruited to the plasma membrane via multivalent interactions with anionic lipids. The CLIC motifs in Bem1 provide the strongest affinity for anionic lipids at this step. 2 Upon their recruitment to anionic lipids, the Bem1 CLIC motifs influence the membrane environment by increasing acyl chain order and rigidifying the local membrane environment.

Previous studies identified an important role for the anionic lipid PS in the anisotropic plasma membrane recruitment of Cdc42 and its regulator Bem1, although the underlying mechanism was unknown ^21^. While the PX domain of Bem1 interacts with PS *in vitro*, mutation of the relevant cationic residues in the Bem1 PX domain did not result in a phenotype unless additional pathways that guide polarity were also inactivated *in vivo* ^15, 17^. It was therefore unknown if additional lipid binding sites existed in Bem1. We demonstrate that Bem1 does indeed employ a second anionic lipid region composed of CLIC motifs, which together with the PX domain, mediates the robust interaction of Bem1 with negatively charged lipids.

Five lines of evidence support the involvement of the CLIC motifs in the anionic lipid targeting of Bem1. First, mutation of the cationic residues that constitute the motifs dramatically reduced the affinity of Bem1 for these lipid species in a reconstituted system. Second, a CLIC motifs construct was sufficient to interact with liposomes mimicking plasma membrane lipid composition. Third, appending a single *CLIC* motif to a *cdc42* mutant that is defective in anionic lipid recruitment is sufficient to restore viability to this otherwise lethal mutant *in vivo*. Fourth, mutation of the *CLIC* motifs reduces the localization of Bem1 to the cell pole at a population and at a single cell level *in vivo*. Finally, as discussed below, solid-state NMR data indicate a specific interaction between Bem1 and anionic lipids.

Anionic lipids recruit Bem1 and this interaction in turn induces ordering of the lipid acyl chain backbone in a PS-PI4P-dependent manner, increasing membrane rigidity. Upon recruitment of Bem1 to anionic lipids, the interaction of the CLIC motifs with these lipids decreases membrane fluidity, likely reducing the diffusion of Cdc42 GTPase components locally. In eukaryotes, diverse Ras-family GTPases display heterogeneous diffusion on the plasma membrane, where active GTPases and other signaling proteins have been imaged in discrete sub-diffraction limited ensembles, also referred to as nanoclusters ^24, 25, 47–52^. Cdc42 is organized in nanoclusters that are larger at the cell pole ^22^, where Bem1 contributes to GTPase activation ^31^. These larger nanoclusters at the pole require Bem1 and phosphatidylserine. Indeed, exogenous addition of PS is sufficient to induce the organization of Cdc42 into large nanoclusters, but only if Bem1 is present ^22^. We therefore propose that Bem1-dependent Cdc42 nanoclustering via PS is likely to be mediated by the CLIC motifs in Bem1. The increased membrane rigidity generated by CLIC-like sequences in other proteins, combined with the potential of PS to span the two leaflets of the lipid bilayer ^23^, may constitute critical ingredients for nanocluster-mediated signaling on the plasma membrane.

Taking advantage of the newly identified CLIC motifs as a starting point, we addressed more generally the mechanistic basis of GEF-scaffold anionic lipid targeting. We identified multivalency in the protein-lipid interactions as a critical constituent of avid GTPase module targeting to the plasma membrane. Intuitively, multivalency as a means of avid protein targeting is appealing, since multiple juxtaposed ligand binding sites in a target have the potential to confer multiplicative rather than additive affinity ^53^. Consistently, a previous study demonstrated that the electrostatic-based interaction of N-WASP with PIP_2_ is multivalent, and that this contributes to both the cooperativity of this protein-lipid interaction and to the ultrasensitive, switch-like kinetics of actin polymerization ^54^ Since polarity establishment is also a switch-like phenomenon, it is feasible that cooperativity in the Bem1-lipid interaction may contribute to these properties. Multivalent protein-lipid interactions also underlie Par polarity complex localization to anionic lipids at the cortex, where basic-hydrophobic domains resemble the CLIC motifs that we identify in Bem1 ^55^. Similarly, multivalent protein-lipid interactions play a role in the recruitment of dynamin, EEA1, retromer and ESCRT-III complexes to membranes ^56–59^. Future experiments examining the degree of lipid penetration by the membrane targeting signals in these proteins are warranted to understand whether they too change the local membrane environment.

In the case of the Cdc42 module in budding yeast, additional peripheral membrane proteins associated with Cdc24 and Bem1 are likely to contribute additional multivalent effects. For example, both Boi1 and Boi2, which interact with Bem1 and Cdc24, each contain a PH domain, as do Cdc42 GAPs ^37, 60, 61^. These proteins may increase the avidity of the Cdc42 GTPase module for PS-PI4P further, or, if their affinity for other lipid species is sufficient, they may contribute additional lipid-specific targeting functions to the GTPase module.

## Methods

### Plasmid construction

Bem1 expression plasmids were generated using a modified pGEX6P-2 backbone in which the BamHI site in the multiple cloning site was changed to NdeI. Full-length Bem1 and the truncated proteins were amplified by PCR, introducing NdeI and XhoI restriction sites and cloned into the modified pGEX6P-2 vector to generate pDM256, pDM548, pDM514, pDM516 and pDM577, respectively. The **clic** mutants *(*clic*-1, *clic*-2, *clic*-3, *clic*-14A* and **clic*-14E)* were synthesized with NdeI and BamHI restriction sites (Bio-Basic, Markham, Canada). The NdeI/BamHI fragments were cloned into the modified pGEX6P-2 vector to generate pDM602 *(*clic*-1)*, pDM604 *(*clic*-2)*, pDM599 *(*clic*-3)*, pDM600 *(c/¡c-14A)* and pDM890 *(c/¡c-14E)*.

*BEM1, bem1 *clic*-14E* and *bem1 *clic*-14E px* constructs were cloned into a yeast integrating plasmid (pRS306) containing 0.4 Kb upstream of the *BEM1* start codon and 0.143 Kb downstream of the stop codon. The *BEM1* coding sequence and mutants were ligated as XhoI-EagI fragments, generating pDM865, pDM906 and pDM947, respectively.

Supplementary Table 1 contains a list of the plasmids used in this study.

### Yeast strains and growth conditions

The *cho1Δ* strains were generated by replacing the *CHO1* gene with kanMX6– or hphNT1-selectable markers ^62, 63^. For experiments employing the *cho1* mutant, minimal medium supplement with 1 μM choline was used, except where noted. *cho1Δ* strains were routinely tested to ensure choline auxotrophy ^64^

The AID-stt4 strains were generated as follows: pDM589 was digested using PmeI to release *TIR1* for integration at *LEU2*. Next, pDM585 was used to generate *pKan-pCUP1-9xMyc-AID-stt4* for Stt4 N-terminal tagging by homologous recombination. Transformants were tested for auxin sensitivity and verified by pcr and DNA sequencing.

The *bem1 px* mutant (K338M, K348A, R349A & R369A) was generated by directed mutagenesis of pDM256. The *bem1 px* coding sequence was then amplified by pcr and transformed into a *bem1Δ::CaURA3* strain (DMY2179). Transformants were selected for loss of the *URA3* marker on 5-FOA and integration of the *bem1 px* mutant was verified by PCR and sequencing, yielding DMY2199.

*BEM1, bem1 clic-14E* and *bem1 clic-14E px* strains were generated using the pop-in-pop-out strategy ^65^. In the first pop-in step, pDM865, pDM906 and pDM947 were linearized by digestion with an enzyme recognizing a restriction site within the *BEM1* or *bem1* ORF, then transformed into DMY2105 and selected on SC-URA for recombination at the *BEM1* locus. In the pop-out step, homologous recombination between the wild type *BEM1* and juxtaposed *bem1* mutant occurs randomly, generating some transformants in which the wild type or mutant *bem1* sequence is present at the *BEM1* locus. After counter selection against *URA3* on 5FOA media, transformants containing untagged *BEM1* or *bem1* at the genomic locus were identified by PCR and DNA sequencing.

The *cdc24 K513A-3xHA* mutant was generated by directed mutagenesis of pDM032, generating pDM737. The wild type or mutant *cdc24 K513A-3xHA* were then integrated at the endogenous *CDC24* locus and checked by PCR, sequencing and western blotting using anti-HA antibodies.

Supplementary Table 2 contains a list of the yeast strains used in this study.

### Protein expression and purification

GST-Bem1, Cdc24-6xHis and derivative mutants were expressed and purified from Bl21-CodonPlus (DE3) cells, essentially as previously described ^31^. Briefly, cells were grown in terrific broth at 37 ºC until an OD600_nm_ ∼3. Expression was induced by the addition of IPTG to 0.3 mM for Bem1 and 0.8 mM for Cdc24, after which cells were grown overnight at 18 ºC. Cells were then harvested and flash frozen in liquid nitrogen. The cell pellets were subsequently ground to a fine powder in a chilled coffee grinder.

For purification of Cdc24-6xHis, room temperature lysis buffer (50 mM Tris-HCL (pH = 8.0), 1 M NaCl, 5 mM imidazole, 5% glycerol, 0.1% tween) supplemented with EDTA-free Protease inhibitor cocktail (Roche, Basel, Switzerland) and 1 mM freshly prepared PMSF was added to the chilled bacterial powder. After sonication on ice, the lysate was centrifuged at 70,000x g for 1 hour and the supernatant was loaded on a Ni2+-IMAC column. Beads were washed with 50 mM Tris-HCL (pH = 8.0), 1 M NaCl, 20 mM imidazole, 5% glycerol, 0.1% tween and Cdc24-6xHis was eluted with 20 mM Tris-HCL (pH = 8.0), 300 mM NaCl, 500 mM imidazole. Cdc24-6xHis was extensively dialysed (50 mM Tris-HCL (pH = 8.0), 150 mM NaCl) then flash frozen in liquid nitrogen for storage. The same protocol was used to purify bem1 CLICs-6xHis, except that the lysis buffer was modified (50 mM Tris-HCL (pH = 7.5), 1 M NaCl, 5 mM imidazole, 5% glycerol, 0.5% tween). The protein was analyzed on a 16% Tris-tricine gel ^66^.

For GST-Bem1 purification, a modified lysis buffer was used (50 mM Tris-HCL (pH = 7.5), 1 M NaCl, 0.1% Tween-20 and 5 mM DTT, EDTA-free Protease inhibitor cocktail and 1 mM freshly prepared PMSF). The lysate was sonicated, centrifuged as above and the supernatant was added to glutathione agarose beads for 2 hours in batch. After extensive washing (50 mM Tris-HCL (pH = 7.5), 250 mM KCl, 0.05% Tween-20 and 0.5 mM DTT), the beads were equilibrated in 3C protease buffer (50 mM Hepes (pH = 7.6), 250 mM KCl, 0.05% Tween-20 and 0.5 mM DTT). The GST tag was digested directly on the glutathione agarose using the same buffer, supplemented with rhinovirus 3C protease. The flow-through, containing untagged Bem1, was dialysed extensively in 50 mM Tris-HCL (pH = 7.5), 150 mM NaCl then flash frozen in liquid nitrogen for storage.

Cdc42 lacking the C-terminal CAAX sequence was tagged with 10xHis and expressed and purified from Bl21-CodonPlus (DE3) cells. Room temperature lysis buffer (50 mM Tris-HCL (pH = 7.5), 1 M NaCl, 25 mM imidazole), supplemented with EDTA-free Protease inhibitor cocktail (Roche, Basel, Switzerland) and 1 mM freshly prepared PMSF was added to bacterial powder. The lysate was stirred, sonicated and centrifuged as described above, and the supernatant was loaded onto a Ni2+-IMAC column. The column was washed in 50 mM Tris-HCL (pH = 7.5), 1 M NaCl, 25 mM imidazole, and Cdc42-10xHis was eluted in 20 mM Tris-HCL (pH = 7.5), 300 mM NaCl and 250 mM imidazole. To obtain nucleotide-free Cdc42, the protein was dialysed in 20 mM Tris-HCL (pH = 7.5), 150 mM NaCl, 5% glycerol supplemented with 25mM EDTA, then dialyzed extensively (20 mM Tris-HCL (pH = 7.5), 150 mM NaCl, 5% glycerol). Samples were flash frozen in liquid nitrogen for storage.

### Liposome preparation

Liposomes were prepared freshly from lipid stocks (Avanti Polar Lipids Inc., Alabaster, USA). The origin and composition of the lipids is provided in Supplemental Table 3. Lipids dissolved in chloroform were lyophilized for 15 minutes at 45 °C to evaporate the chloroform. Lipids were washed in 50 μL ultra-pure water and lyophilized until dry. Lipids were the resuspended in 20 mM Tris (pH = 7.5), 150 mM NaCl to have a final lipid concentration of 2 mM. After 6 cycles of freeze-thaw in liquid nitrogen and at 45°C in a water bath, liposomes were sonicated for 15 minutes in a bath sonicator. This method yielded monodisperse preparations of ∼100 nm diameter liposomes, as assessed by dynamic light scattering.

### Liposome floatation assays

The final concentration of liposomes was 0.5 mM in floatation experiments, while protein was 2 μM unless indicated differently. Liposomes were mixed with buffer alone (20 mM Tris (pH = 7.5), 150 mM NaCl), or with protein, in a final volume of 150 μL in a 500 μL polycarbonate ultracentrifuge tube. The mixtures were incubated at room temperature for 30 minutes. 100μL of a 75 % sucrose solution was mixed with the protein-liposome mixture to give a final sucrose concentration of 30%, which was gently overlaid with 200 μL of 25% sucrose. Finally, 50 μL of 20 mM Tris (pH = 7.5), 150 mM NaCl was overlaid to give a final volume of 500 μL. Tubes were centrifuged for 1 hour at 23°C at 120,000x g. 100 μL of supernatant and 200 μL of pellet were precipitated in 10% Tri-Chloroacetic Acid. The pellet was resuspended in 15 μL of SDS-PAGE sample buffer (65 mM Tris-HCl (pH = 6.8), 2% SDS, 10% glycerol, 5% β-mercapto ethanol, 100 mM ß-glycerophosphate, 50 mM sodium fluoride), boiled for 5 minutes, analyzed by SDS-PAGE and stained with Coomassie brilliant blue R250. The intensity of the protein bands in the supernatant and pellet were analyzed using a Bio-Rad Gel Doc system running Image Lab software.

The percentage of floating protein was calculated using the equation: {Supernatant ^band intensity^ / (Supernatant ^band intensity^ + Pellet ^band intensity^)} * 100

### Solid-state NMR spectroscopy

Liposomes containing DMPC-d54, DMPS and brain PI(4)P were prepared by mixing powders in organic solvent (chloroform/methanol, 2:1) in the presence or absence of the Bem1 CLIC motifs (amino acids 1–72) and adjusting the lipid/protein ratio (25:1). Solvent was evaporated under a flow of N_2_ to obtain a thin lipid film. Lipids were rehydrated with ultrapure water before lyophilisation. The lipid powder was hydrated with an appropriate amount of deuterium-depleted water (80 % hydration ratio) and homogenized by three cycles of vortexing, freezing (liquid nitrogen, −196°C, 1 min) and thawing (40°C in a water bath, 10 min). This protocol generated a milky suspension of micrometer-sized multilamellar vesicles.

^2^H NMR spectroscopy experiments were performed using a Bruker Avance II 500 MHz WB (11.75 T) spectrometer. ^2^H NMR spectroscopy experiments on ^2^H-labelled DMPC were performed at 76 MHz with a phase-cycled quadrupolar echo pulse sequence (90°x-t-90°y-t-acq). Acquisition parameters were as follows: spectral window of 500 kHz for ^2^H NMR spectroscopy, p/2 pulse width of 3.90 ms for ^2^H, interpulse delays (t) were of 40 ms, recycled delays of 2 s for ^2^H; 3000 scans were used for ^2^H NMR spectroscopy. Spectra were processed using a Lorentzian line broadening of 300 Hz for ^2^H NMR spectra before Fourier transformation from the top of the echo. Samples were equilibrated for 30 min at a given temperature before data acquisition. All spectra were processed and analyzed using Bruker Topspin 3.2 software. Spectral moments were calculated for each temperature using the NMR Depaker 1.0rc1 software [Copyright (C) 2009 Sébastien Buchoux]. Orientational order parameters (S_CD_) were calculated from experimental quadrupolar splittings (Dn _Q_) after spectra simulations according to equation 1 :

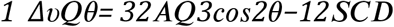

in which A_Q_, the quadrupolar coupling constant for methyl moieties is 167 kHz, and q is the angle between the magnetic field and the bilayer normal.

### Western blotting

Samples for Western blotting were prepared by collecting 1 OD600_nm_ of mid-logarithmic phase cells, adding glass beads and flash freezing in liquid nitrogen ^37^. Samples were vigorously agitated in 60 μL SDS sample buffer supplemented with 1 mM fresh PMSF. Samples were immediately boiled and analyzed by SDS-PAGE, Western blotting and probed with appropriate antibodies.

### Imaging

Cells were imaged using a wide-field inverted microscope (Zeiss Axiovert 200M) with a 100× objective (oil, numerical aperture [NA] 1.4, plan Apo), and an electron-multiplying charge-coupled device (EMCCD) camera (Evolve; Photometrics, Tuscon, AZ). MetaMorph 7.7 software (Molecular Devices, Sunnyvale, CA) was used for image acquisition and analysis. Filter sets LF488-B-000 (FFO2-482/18, FFO1-525/45, Di01-R488 [exciter, emitter, dichroic]) and LF561-A-000 (FFO2-561/14, FFO1-609/54, Di01-R561) were used to sequentially image cells expressing Bem1-GFP and Cdc24-mCherry ^67^.

### Image analysis

Deconvolution was performed for visualization, where indicated, using a plug-in running within Metamorph software ^68^. All Images were analyzed and processed using ImageJ software on raw data, not on deconvolved images.

To calculate the enrichment of PI4P at the plasma membrane, the integrated intensity (II) and area (A) of the entire cell (E) and the cytosol (C) were determined for each cell after background subtraction. Next, the mean gray value of the plasma membrane (MGVP) for each cell was determined as follows: MGVP = (IIE-IIC)/(AE-AC)). The values were plotted using GraphPad Prism software.

To calculate the enrichment of Bem1 at the pole, the mean gray value of the pole (MGVP) and the cell (MGVC) were determined for each cell using an empirically determined threshold value that enabled the cell pole to be identified. Next, the MGVP was normalised as follows: normalised mean gray value of the pole = (MGVP-MGVC)/MGVC). The values were plotted using GraphPad Prism software.

### Cdc24 GEF assay

Förster resonance energy transfer (FRET) between Cdc42 and N-methylanthraniloyl-GTP (mant-GTP) was measured to monitor the Cdc24-mediated GDP to mant-GTP exchange reaction on Cdc42 in real time. Trp97 of Cdc42, which is in close proximity to the GTP binding site, was excited using 280 nm wavelength light using a 5 nm bandwidth. The FRET signal was detected at the emission peak of mant-GTP, at 440 nm using an 8 nm bandwidth. All fluorescence measurements were performed at 27°C on a Tecan Infinite M1000PR0 plate reader (Tecan Group, Männedorf, Germany) in 384-well, non-binding microplates (Greiner Bio-One, Courtaboef, France), in a 10 μl reaction volume. The final buffer conditions were 20 mM Tris-HCl (pH = 8.0), 150 mM NaCl, 1 mM DTT, 5 mM MgCl2, 100 nM mant-GTP, supplemented with 100 μM GMP-PNP nucleotide. Cdc24 was used at a final concentration of 60 nM after 30 min room temperature pre-incubation with Bem1 at 5 μM. The reaction was initiated by adding Cdc42 to 9 μM final concentration and exchange was monitored for at least 2000 s with 15 s intervals. For each sample, a mock reaction was used in the absence of GDP-loaded Cdc42 to normalise for bleaching and to subtract possible sources of background noise such as Cdc24-mant-GTP interaction. The intrinsic GDP to mant-GTP exchange rate of Cdc42 was determined in the absence of Cdc24.

Fitting of the kinetic trace data was performed in GraphPad Prism using a single exponential equation and the observed kinetic rate constants were compared.

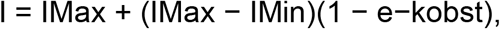

where I is the fluorescence intensity change, IMax is the maximal fluorescence intensity, IMin is the minimal fluorescence intensity, kobs is the observed kinetic rate constant and t is the time in seconds.

### Cdc24 affinity for anionic lipids in the presence of Bem1

The affinity of Cdc24 for anionic lipids in the presence of Bem1 was estimated using nonlinear regression analysis ^69^. In the analysis, we assume an approximate initial K_d_ between Cdc24 and anionic lipids of 1 μM ^14^. A nonlinear regression fit with equation 2 was used to obtain the corresponding K_d_ for the Cdc24 interaction with anionic lipids in the presence of Bem1:

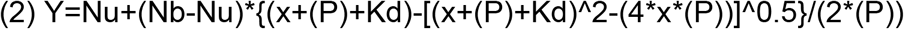

where Y is the percentage of Cdc24 found in the supernatant at the lipid concentration P. P was chosen based on the following: first, around 30% of Cdc24 interacted with anionic liposomes (see Figure 5B). Second, the PH domain is assumed to interact with 3-5 lipid molecules ^70^. In the floatation experiments involving Bem1 and Cdc24 shown in Figure 5B, C and D, 1 μM of Cdc24 was used, so P = [Cdc24]*(30%)*3 lipid molecules = 0.9 μM lipids that are estimated to interact with Cdc24. In the equation, Nu and Nb are the maximum unbound and bound percentage of Cdc24, respectively, and x is the Bem1 concentration.

### Single Particle Tracking Photoactivation Localization Microscopy (sptPALM)

Live cells were imaged using a widefield, inverted microscope (Axiovert 200M; Carl Zeiss, Marly le Roi, France) equipped with a 100X TIRFM objective (oil, NA 1.46, plan Apo), iLas^2^ TIRF system (Roper Scientific) and an EMCCD camera (Evolve; Photometrics, Tuscon, Arizona). The imaging system was maintained at a constant temperature of 25°C using a custom designed incubator (Box and Cube, Life Imaging System, Basel, Switzerland). MetaMorph 7.7 software (Molecular Devices, Sunnyvale, USA) was used for image acquisition and analysis.

For *in vivo* imaging, cells expressing mEOS-Cdc42 were grown to midlog phase and imaged at 25°C. Coverslips (High precision 18 × 18 mm, 1.5 H, Marienfeld, Lauda-Königshofen, Germany) were washed overnight in a solution of 1M HCl and 1M HNO3 then rinsed three times the next day in ultrapure water. After a 30-minute incubation in water, then 30 minutes in ethanol, the coverslips were dried and used for imaging. Imaging was performed in a highly oblique illumination (HiLo) mode. mEOS-Cdc42 cells were imaged using a 561 nm laser with additional continuous photoconversion using a 405 nm laser. The 405 nm laser was maintained at low power (0.3–1 μW) for adequate separation of stochastically converted molecules. The iLas^2^ system was used in arc mode for live imaging and ellipse mode for fixed samples. These settings set the pattern of rotation of the lasers on the back focal plane of the TIRF objective. The fluorescence was collected on the EMCCD camera after passing through a combination of dichroic and emission filters (D101-R561 and F39-617 respectively; Chroma, Bellows Falls, VT). Images were acquired in streaming mode at 50 Hz (20 ms exposure time). During *in vivo* imaging, 16,000 to 20,000 images were collected for each cell. Multicolour fluorescent 100 nm beads (Tetraspeck, Invitrogen) were used as fiduciary markers in all super-resolution imaging experiments to register long-term acquisitions for lateral drift correction.

For fixed-cell imaging, cells were grown to log phase (OD_600 nm_ of <0.8) and fixed with 3.7% formaldehyde and 0.2% glutaraldehyde for 10 minutes. After washing in PBS three times, cells were resuspended in PBS and directly used for imaging. Image acquisition of fixed cells was performed using the same protocol as for living cells, as described above. 32,000 to 40,000 images were acquired per cell, at which point the pool of photoconvertible single molecules was completely depleted.

### Single particle localization, tracking and nanocluster detection by Voronoï Tesselation

Image stacks collected for each sptPALM experiment were analyzed using a custom-written software operating as a plugin within MetaMorph software, PalmTracer, to compute single molecule localizations and dynamics. Diffusion coefficients obtained for each strain are listed in Table 1. Single molecules were localized in each image frame and tracked over time using wavelet segmentation and simulated annealing algorithms ^22^. The sptPALM image resolution, defined as FWHM = 2.3 x the pointing accuracy, was estimated to 48 nm. The pointing accuracy, measured to be 20.86 nm, was computed from the acquisition of mEOS-Cdc42 in fixed cells by bidimensional Gaussian fitting of the spatial distribution of 80 single molecules localized for more than 20 consecutive time points. Tracking data and subsequent MSDs were generated from the membrane-bound population of mEOS-Cdc42. Proteins in the freely diffusing cytosolic pool of mEOS-Cdc42 were not tracked in these experiments because cytosolic diffusion is much higher than diffusion in a membrane environment, and would not be localized and tracked with 20 ms exposure time.

In our observations, all MSDs have a quasi-linear dependence at short times, enabling computation of the instantaneous diffusion coefficient (D) per molecule by linear regression on the first four points of the MSD of all trajectories that are longer than 6 consecutive frames.

Cdc42 nanoclusters were quantified from the reconstructed superresolution images of fixed cells using SR-Tesseler analysis ^22^. This software is based on Voronoï tessellation, wherein single molecule localizations are treated as seeds around which polygons are assembled. In our analysis, we defined regions of interest (ROI) as the pole or non-pole of the cell after visual inspection of the widefield 491 nm image acquired at the outset of the experiment. The surface area of the polygon drawn around the detected single molecule is proportional to the local molecular density, such that the area of the polygon decreases as the local density of single molecule localizations increases. PALM images were corrected for single-molecule blinking within the SR-Tesseler software ^22^. This takes into account mEOS photophysics, and a pointing accuracy of 20 nm as a radius of search, which would otherwise overestimate the number of single molecule detections. After blinking correction, nanoclusters were defined as those areas containing a minimum of five localizations at a local density that was at least two-fold higher than the average density within the selected ROI. Nanocluster characteristics including diameter, area and the number of localizations were exported from SR-Tesseler into Excel (Microsoft) for further statistical analysis.

### Statistical Analysis

The diffusion coefficients were represented as box plots displaying the median as a line and the percentiles (25-75%). Statistical comparisons were made using a non-parametric, two-tailed Mann-Whitney rank sum test. Non-Gaussian distributions of nanocluster sizes were represented by data-points displaying median as a line and the percentiles (25–75%) and also compared using a non-parametric, two-tailed Mann-Whitney rank sum test. Statistical analyses were based on cluster area values calculated by SR-Tesseler. Only areas greater than 2000 nm^2^ were used, corresponding to a diameter of 48 nm, the resolution of our imaging system.

## Acknowledgements

We thank Cameron Mackereth for advice on nonlinear regression analysis and fitting methods. We also thank Bertrand Daignan-Fornier for anti-Ade13 antibody, Helle Ulrich and Douglas Koshland for AID/TIR plasmids and Nelly Savarit, Melanie Sevrin and Laure Bataille for performing preliminary liposome floatation experiments. Anne Royou and Andrew Weatherall are acknowledged for their continued support. We thank Jean-Louis Mergny for the use of his fluorescence plate reader. This work received funding from the University of Bordeaux through the Synthetic Biology in Bordeaux (SB2) Program and a Doctoral School Fellowship to JM. This work was also funded by the CNRS, ANR through Program Blanc grant ANR-13-BSV2-0015-01 and ANR-14-CE09-0020-01, the Regional Council of Aquitaine, the European Research Council (ERC-2015-StG GA no. 639020 to A.L.) and the IdEx Bordeaux (Chaire d’Installation to B.H., ANR-10-IDEX-03-02).

## Author contributions

DMcC conceived the study. All authors designed experiments and analyzed the data. JM and AML performed the experiments, with the exception of NMR spectroscopy, which was performed by DM and analyzed by AL and BH. DMcC wrote the manuscript.

---

**Supplementary Fig. 1.**
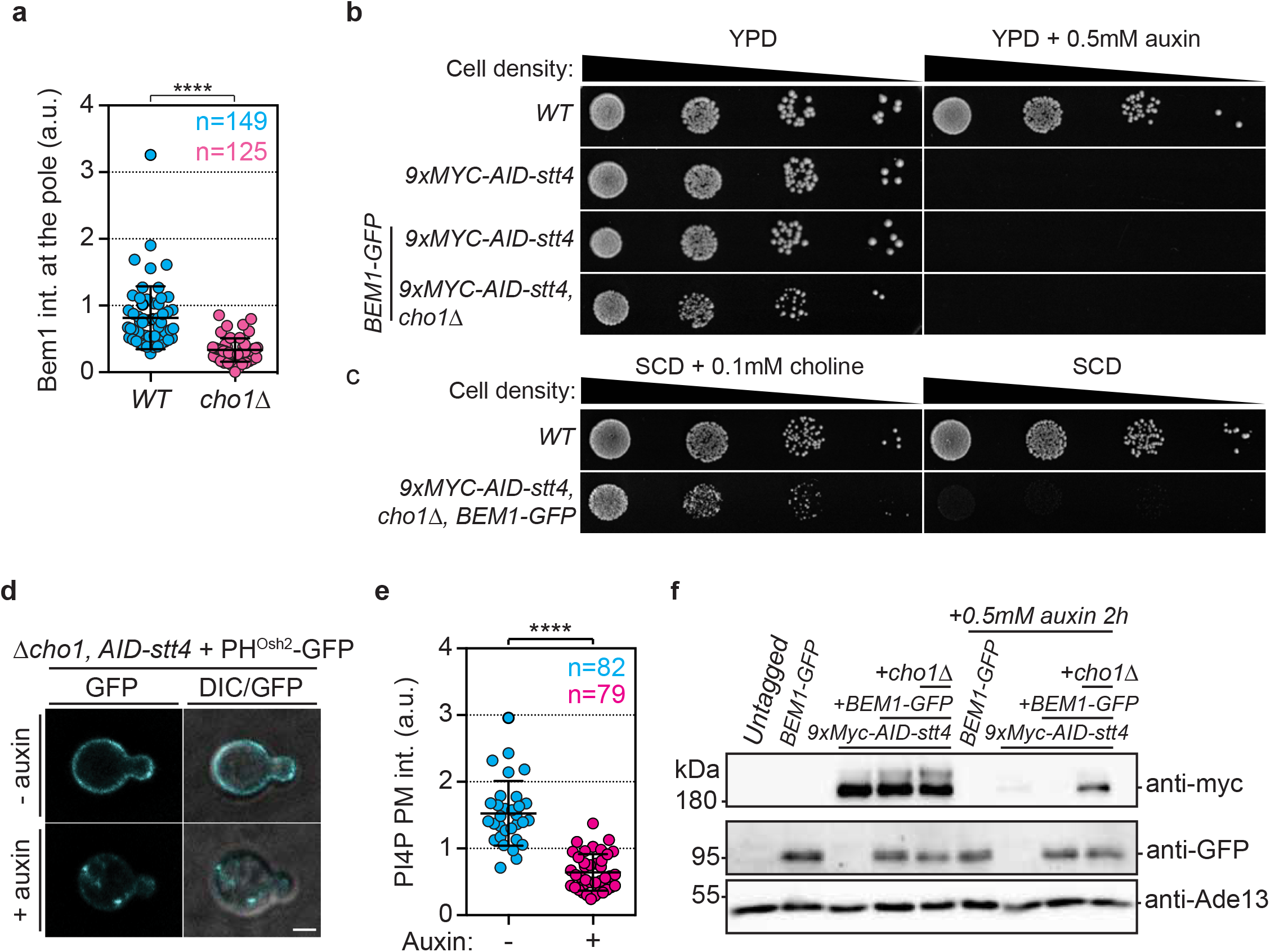
**a**, Scatter dot plot showing the level of Bem1-GFP signal at the cell pole. (n> 100 cells observed from 3 experiments). Bars correspond to mean and SD. Mann-Whitney tests were performed where confidence is ****p<0.0001). **b**, Ten-fold serial dilutions and subsequent colony formation of cells of the indicated genotype. Cells were grown at 30°C for two days on YPD plates with or without 0.5 mM auxin. **c**, Ten-fold serial dilutions and subsequent colony formation of cells of the indicated genotype. Cells were grown at 30°C for three days on SCD plates with or without 0.1 mM choline. **d**, Representative images of a GFP-tagged PI4P probe signal (GFP-2xPH^Osh2^) (cyan) in *cho1Δ, 9xMyc-AID-stt4* cells after 2h treatment with or without 0.5 mM auxin. Merged DIC-fluorescence images are also shown. Images show average intensity projections of deconvolved z-stacks. **e**, Scatter dot plot of PI4P levels in *cho1Δ 9xMyc-AID-stt4* cells after 2h treatment with or without 0.5 mM auxin (n>80 cells in 3 independent experiments). Bars correspond to mean and SD. Mann-Whitney tests were performed where confidence is ****p<0.0001. **f**, Western blots probed with anti-Myc antibody (top panel) to detect AID-stt4, anti-GFP antibody (middle panel) to detect Bem1 or anti-Ade13 antibody (bottom panel) as a loading control. Note that Bem1 levels in *cho1Δ, 9xMyc-AID-stt4* cells treated with auxin for 2h remain similar to the wild type.

**Supplementary Fig. 2.**
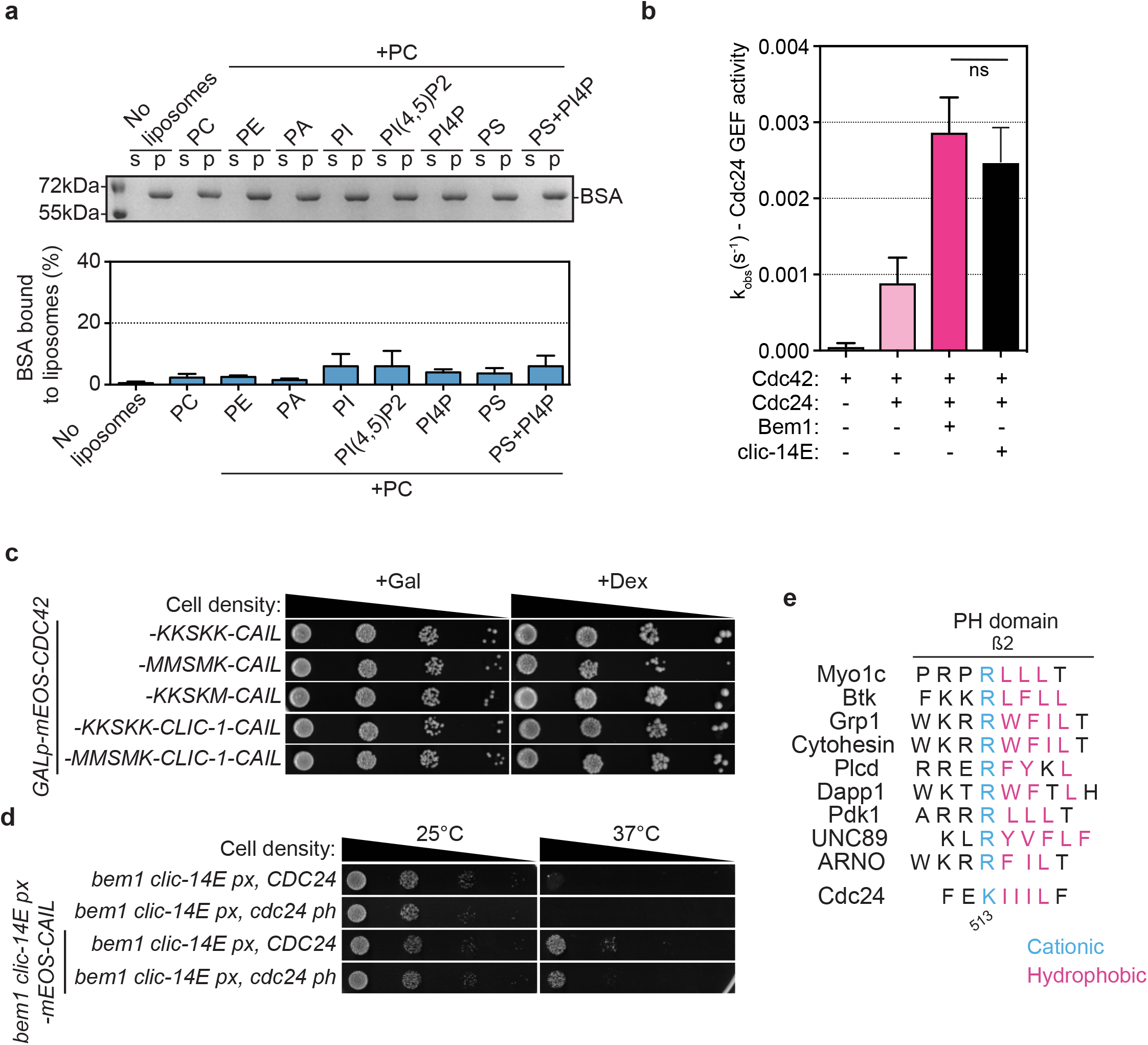
**a**, Upper panel. SDS-PAGE stained with Coomassie blue displaying BSA. Lower panel. Quantification of the percentage of BSA associated with liposomes containing the lipids indicated. Error bars display SD (n=3 experimental replicates). **b**, Observed kinetic rate constants of GEF loading of mant-GTP Cdc42. Values were obtained by fitting trace data to a single exponential equation. Error bars show SD. Values were compared using Student t-tests. **c**, and **d**, Ten-fold serial dilutions of the cells indicated were grown at the temperatures displayed for three days. **e**, Alignment of the second β sheet in the PH domain of the proteins indicated. Note how a conserved cationic residue is followed by a hydrophobic patch.

**Supplementary table 1.**
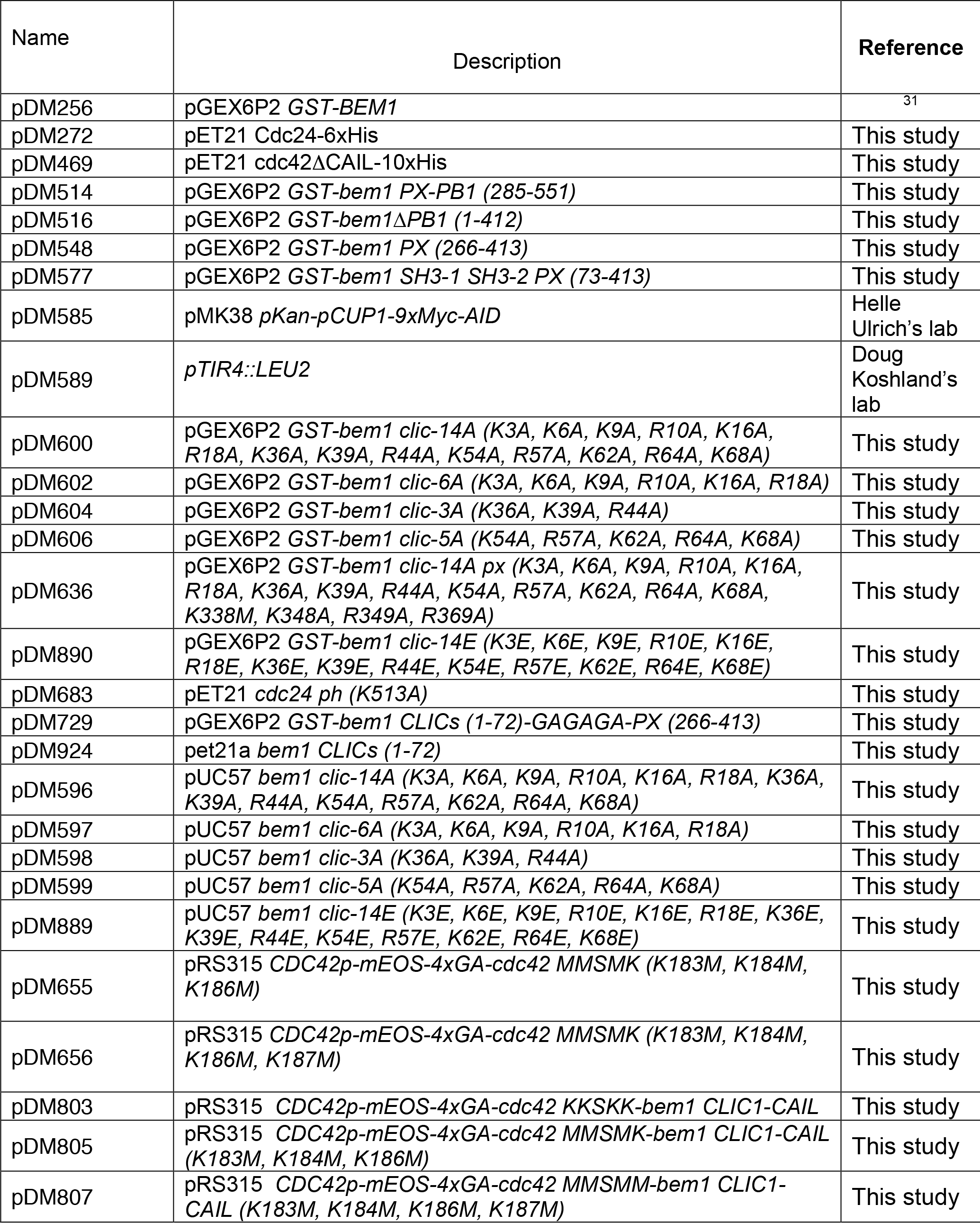
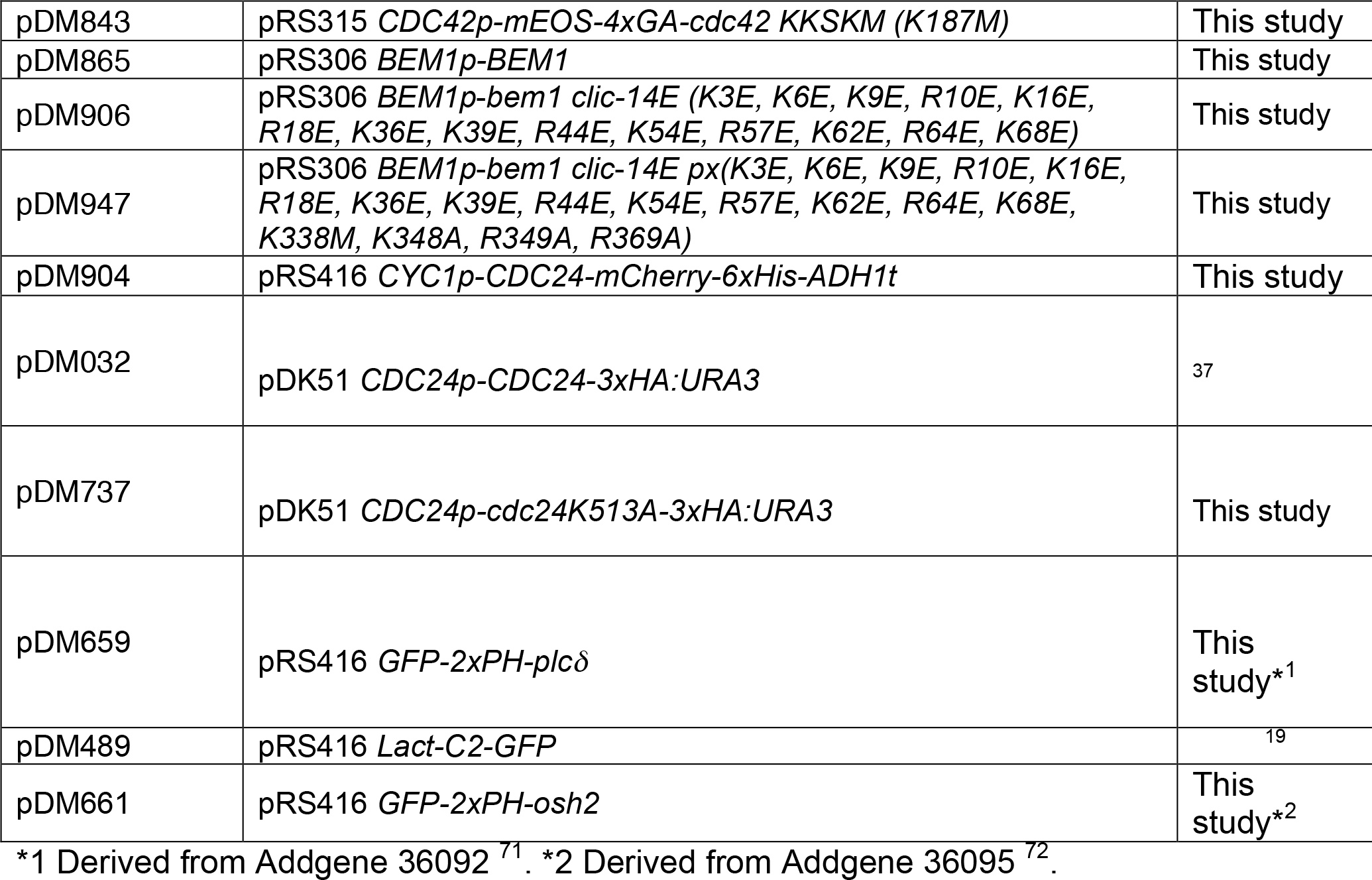
Plasmid constructs.

**Supplementary table 2.**
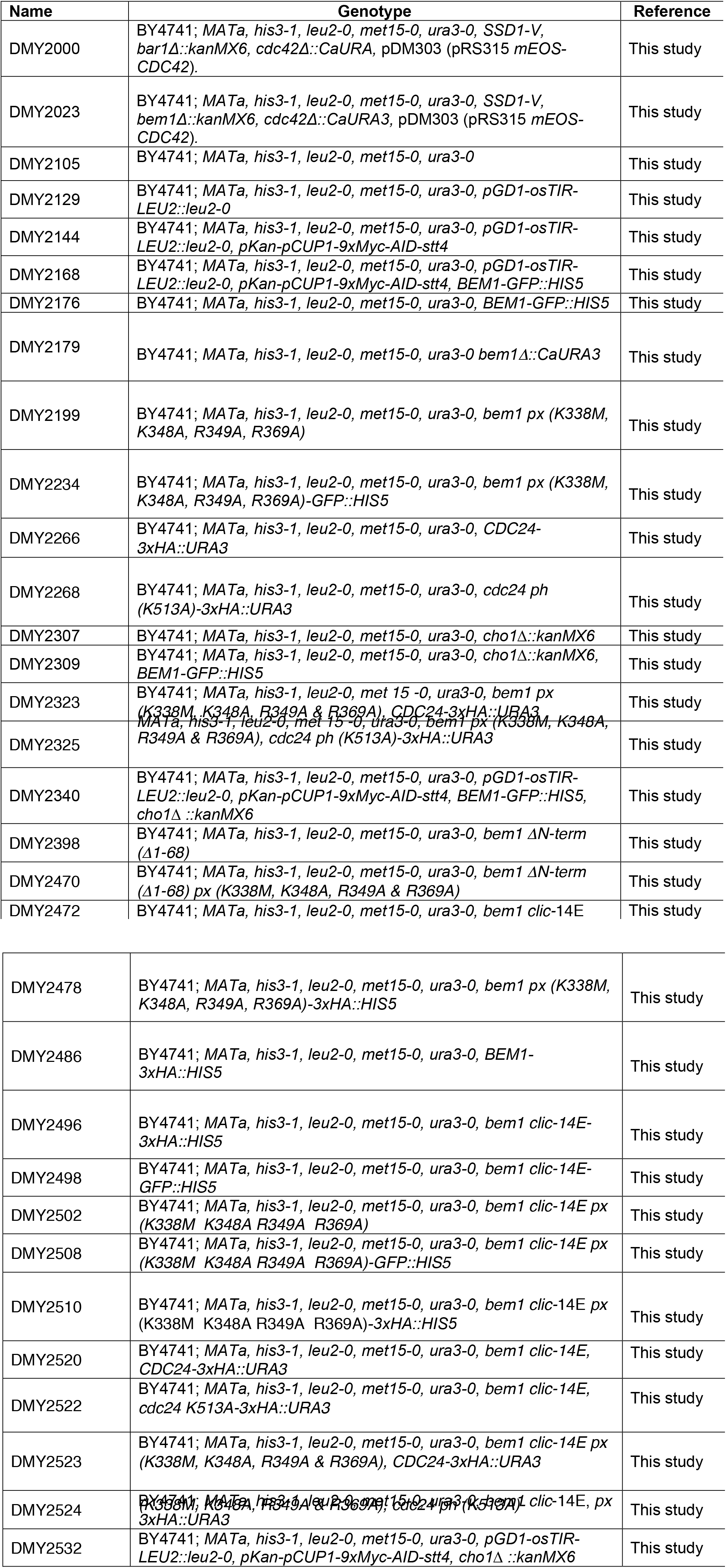
Yeast strains.

**Supplementary table 3.**
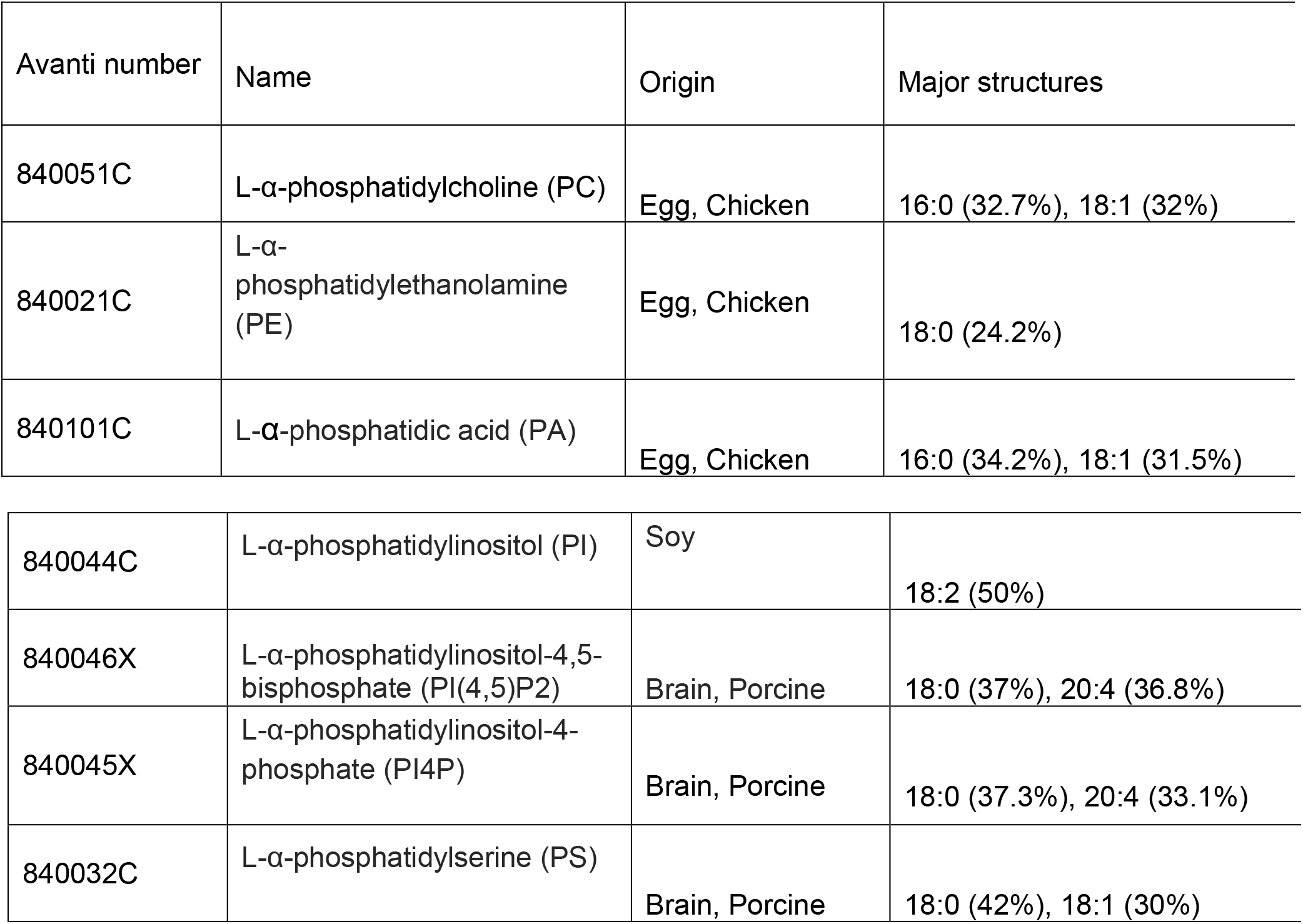
Source of lipids and composition.

